# The type VII secretion system protects *Staphylococcus aureus* against antimicrobial host fatty acids

**DOI:** 10.1101/572172

**Authors:** Arnaud Kengmo Tchoupa, Kate E. Watkins, Rebekah A. Jones, Agnès Kuroki, Mohammad Tauqeer Alam, Sebastien Perrier, Yin Chen, Meera Unnikrishnan

**Affiliations:** Warwick Medical School, University of Warwick, Coventry, United Kingdom; Department of Chemistry, University of Warwick, Coventry, United Kingdom; Faculty of Pharmacy and Pharmaceutical Sciences, Monash University, Parkville, Victoria 3052, Australia; School of Life Sciences, University of Warwick, Coventry, United Kingdom

**Keywords:** *Staphylococcus aureus*, Type VII secretion system, long-chain unsaturated free fatty acids

## Abstract

The *Staphylococcus aureus* type VII secretion system (T7SS) exports several proteins that are pivotal for bacterial virulence. The mechanisms underlying T7SS-mediated staphylococcal survival during infection nevertheless remain unclear. Here we show that the absence of EsxC, a small secreted effector implicated in bacterial persistence, results in cell membrane defects in *S. aureus*. Interestingly, isogenic mutants lacking EsxC, other T7SS effectors EsxA and EsxB, or the membrane-bound ATPase EssC, are more sensitive to killing by the host-derived antimicrobial fatty acid, linoleic acid (LA), compared to the wild-type (WT). LA induces more cell membrane damage in the T7SS mutants compared to the WT. Although WT and mutant strains did not differ in their ability to bind labelled LA, membrane lipid profiles show that T7SS mutants are less able to incorporate LA into their membrane phospholipids. Furthermore, proteomic analyses of WT and mutant cell fractions reveal that, in addition to compromising membranes, T7SS defects induce oxidative stress and hamper their response to LA challenge. Thus, our findings indicate that T7SS is crucial for *S. aureus* membrane integrity and homeostasis, which is critical when bacteria encounter antimicrobial fatty acids.

## Introduction

*Staphylococcus aureus* is a facultative pathogen that can colonize the skin and nares of healthy individuals. The asymptomatic carriage of *S. aureus* is a major risk for subsequent infections (von Eiff *et al*., 2001). *S. aureus* infections, which can be healthcare or community-associated, range from benign impetigo to life-threatening bacteraemia (Tong *et al*., 2015). Clinical management of staphylococcal infections is complicated by the increasing prevalence of multidrug resistant strains (Lee *et al*., 2018).

The success of *S. aureus* as a deadly pathogen is attributed to an array of virulence factors that facilitate host tissue adhesion and immune response evasion (Gordon & Lowy, 2008). One of these virulence factors is the type VII secretion system (T7SS), also known as the ESAT-6 secretion system (ESS). The orthologous ESX-1 system was initially discovered in *Mycobacterium tuberculosis*, where it is essential for bacterial virulence (Conrad *et al*., 2017). T7SSs (T7SSb) are found in both Gram-positive and Gram-negative bacteria, although these systems and their secretion machineries appear to be distinct to their mycobacterial counterparts (Unnikrishnan *et al*., 2017).

In extensively studied strains (COL, RN6390, USA300 and Newman), the T7SS consists of four integral membrane proteins (EsaA, EssA, EssB and EssC), two cytosolic proteins (EsaB and EsaG), five secreted substrates (EsxA, EsxB, EsxC, EsxD and EsaD), and EsaE, which interacts with the T7SS substrates to target them to the secretion apparatus (Cao *et al*., 2016). A peptidoglycan hydrolase, EssH, was reported to mediate T7SS transport across the bacterial cell wall envelope (Bobrovskyy *et al*., 2018).

The molecular architecture of the staphylococcal T7SS has not yet been fully characterized. T7SS integral membrane proteins EsaA, EssA, EssB, and EssC are thought to be the core of the T7 secretion machinery, with EssC being the central membrane transporter (Burts *et al*., 2005, Jager *et al*., 2018, Zoltner *et al*., 2016). Interactions between secreted substrates and co-dependent secretion of substrates have been demonstrated (Anderson *et al*., 2013, Cao *et al*., 2016, Kneuper *et al*., 2014, Ohr *et al*., 2017). A recent study showed that the functional assembly of the T7SS machinery in *S. aureus* is supported by the flotillin homolog FloA, within functional membrane microdomains (Mielich-Suss *et al*., 2017).

The *S. aureus* T7SS is pivotal for bacterial virulence. Indeed, S. *aureus* mutants lacking the entire T7SS (Kneuper *et al*., 2014) or specific T7SS components (EsxA, EssB, EssC, EsxC, EsxB, EsaB, EsaD or EsaE) were consistently shown to be less virulent and/or persistent in various mouse infection models (Anderson *et al*., 2011, Anderson *et al*., 2017, Burts *et al*., 2008, Burts *et al*., 2005, Ishii *et al*., 2014, Lopez *et al*., 2017). EsxA is necessary to delay apoptosis of *S. aureus*-infected epithelial and dendritic cells, while other substrates modulate cytokine production (Anderson *et al*., 2017, Cruciani *et al*., 2017, Korea *et al*., 2014). Although the relevance of T7SS to *S. aureus* is less understood, a role for the toxin-antitoxin pair EsaD (or EssD) and EsaG (or EssI) was recently demonstrated in intraspecies competition (Cao *et al*., 2016, Ohr *et al*., 2017).

Interestingly, *S. aureus* T7SS expression is induced in response to host-specific FAs (Ishii *et al*., 2014, Kenny *et al*., 2009, Lopez *et al*., 2017), although the role of T7SS in bacterial resistance to antimicrobial FAs remains unclear. In this study, we demonstrate that *S. aureus* lacking the T7SS substrate EsxC has a defective cell membrane. Intriguingly, EsxC and other T7SS mutants were more sensitive to an unsaturated FA, linoleic acid (LA), compared to the wild-type (WT). Although there were no differences in binding labelled LA, LA induced a more leaky membrane in the T7SS mutants, and there was less incorporation of LA into membrane phospholipids. Cellular proteomics revealed that in addition to membrane discrepancies, T7SS mutants exhibited different redox and metabolic states, which likely result in a distinct response to LA.

## Results

### The type VII substrate EsxC is present on the staphylococcal surface and affects its composition

EsxC, a small 15-kDa protein secreted by the T7SS, is important for *S. aureus* persistence in mice (Burts *et al*., 2008). However, mechanisms underlying EsxC- or T7SS-mediated bacterial survival are not known. In order to understand the role of EsxC, we generated an isogenic *esxC* mutant as described previously (Bae & Schneewind, 2006), and confirmed the absence of any secondary site mutations by whole genome sequencing. Δ*esxC* had a similar growth rate to the WT USA300 JE2 strain (Fig. S1). EsxC is known to be a secreted effector of T7SS (Burts *et al*., 2008), although it has been consistently detected in cell membrane (CM) fractions of RN6390 and USA300 (Bobrovskyy *et al*., 2018, Kneuper *et al*., 2014). As reported previously, we detected EsxC in the CM fractions of the WT but not in Δ*esxC* (Fig. S2). Moreover, we also detected EsxC in the WT cell wall (CW) fraction (Fig. S2). In a deletion mutant of the membrane-bound major ATPase EssC (a core T7SS component), EsxC was detected in CM and CW preparations even in the absence of EssC (Fig. S2). While we were unable to detect EsxC in the cell-free supernatants by immunoblotting, we detected this protein by secretome analyses. Along with EsxC, other T7SS proteins, EsxA and EsxD, were detected in the WT but not in Δ*esxC or* Δ*essC* supernatants (Table S1).

Given the association of EsxC with the cell membrane, we stained the WT, Δ*esxC* and Δ*essC* mutants with FM-143, a fluorescent cell membrane probe (Pulschen *et al*., 2017). We observed a mild but statistically significant increase in FM-143 staining in the mutants as compared to the WT (Fig. 1A). Increased staining of membranes by FM-143 has been associated previously with membrane blebbing in bacteria (Wood *et al*., 2019).

**Fig. 1.**
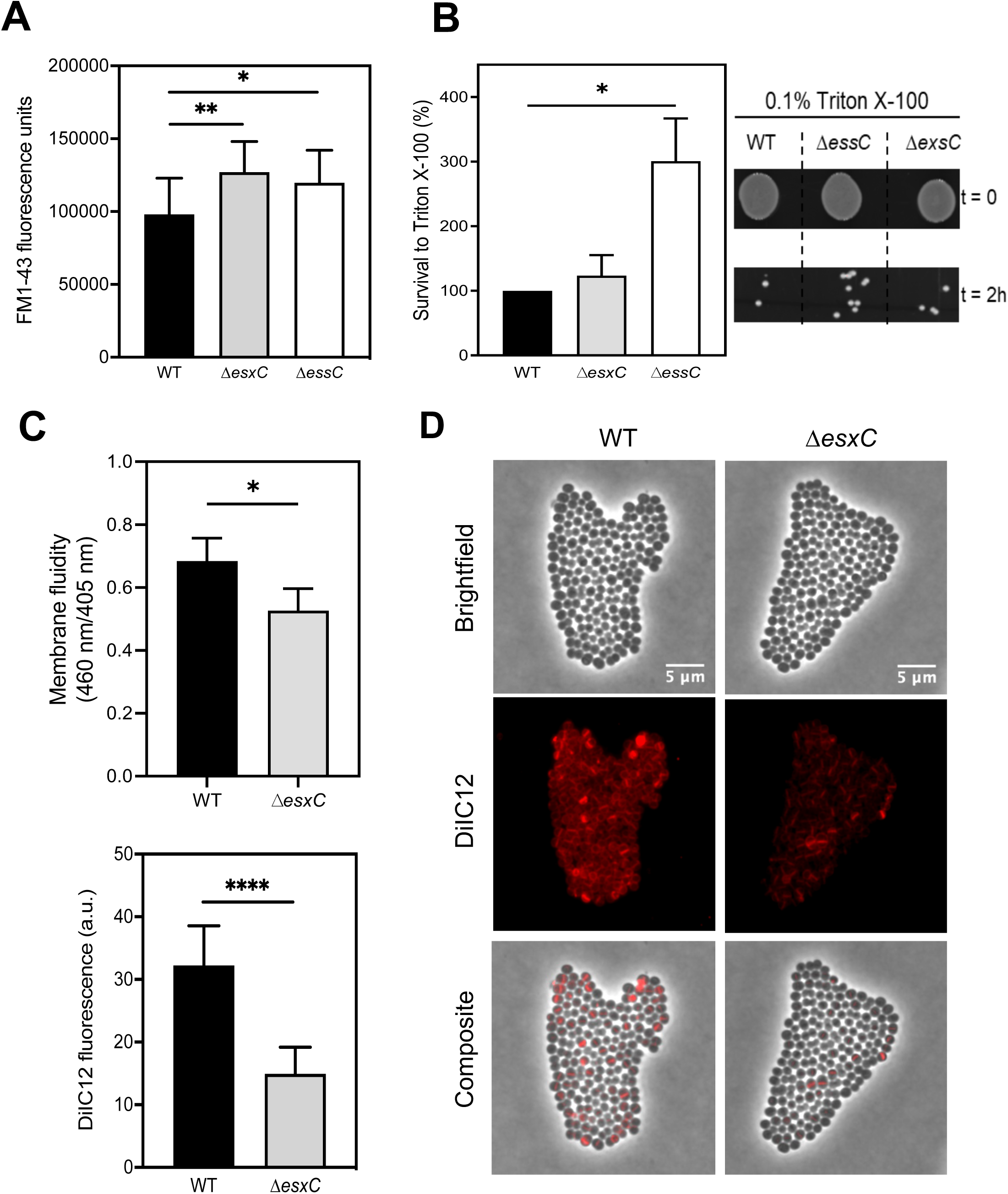
Membrane defects in the absence of EsxC. **A.** FM1-43 fluorescence of *S. aureus* WT, Δ*essC*, and Δ*esxC* as measured with a plate reader. Means ± standard deviation (SD) are shown, *n* = 6; * indicates *P* < 0.05 using a Kruskal-Wallis test with Dunn’s test. **B.** Survival of Δ*esxC* or Δ*essC* after a 2h-treatment with 0.1% Triton X-100 relative to the WT. Means are shown; error bars represent standard deviation (SD). *n* = 4, * indicates *P* < 0.05 using a Kruskal-Wallis test with Dunn’s test. The panel on the right shows a representative picture of bacteria before and after Triton X-100-treatment. **C.** The membrane fluidity of WT and Δ*esxC* as measured with a pyrene decanoic acid staining-based assay. Mean values are shown; error bars represent standard error of the mean (SEM). *n* = 6, * indicates *P* < 0.05 using a two-tailed t-test. **D.** Widefield micrographs of *S. aureus* WT and Δ*esxC* after growth in TSB to OD_600_ of 1.0 in the presence of the lipophilic dye DiIC12. Images are representative of 4 independent experiments. **E.** The DiIC12 fluorescence of 80 bacterial clusters from different fields per strain was quantitated with ImageJ. Means ± standard deviation (SD) are shown, *n* = 4; indicates **** *P <* 0.0001, using a Kruskal-Wallis rank test

To study if the lack of T7SS proteins affected other surface proteins, a surface proteome analysis of the WT and mutants was carried out using trypsin treatment to digest surface proteins (Solis *et al*., 2014). We found that most proteins (172/218) were less abundant in Δ*esxC* compared to the WT (Fig. S3). Interestingly, majority of relatively less abundant proteins (16/20) in the Δ*esxC* surface proteome were predicted or proven to be cytosolic (Table S2). Metabolic (PfKA, FolD, FabI GatB, and ImdH) or stress-related (Asp23 and putative universal stress proteins) proteins were lowest in abundance (log_2_ fold change < - 1.0). The T7SS core component EsaA, which has a prominent extracellular loop (Dreisbach *et al*., 2010), was also ∼50% less abundant in Δ*esxC*, implying that EsxC absence may affect T7SS assembly. Identification of cytosolic proteins during surface proteome analysis could be attributed to non-canonical secretion of cytosolic proteins that bind to bacterial surfaces (Hempel *et al*., 2011, Pasztor *et al*., 2010) or cell lysis during sample preparation (Dreisbach *et al*., 2010, Solis *et al*., 2014, Ventura *et al*., 2010). To study if the T7SS mutants were more resistant to cell lysis, we compared the Triton X-100 lysis of WT, Δ*esxC* and, Δ*essC* in PBS in non-growing conditions. While Δ*esxC* was slightly more resistant to Triton X-100, Δ*essC* was clearly defective in lysis (Fig. 1B), as compared to the WT.

We probed T7SS effects on the membrane further by assessing membrane fluidity of the WT and Δ*esxC* mutant. We used pyrene decanoic acid, an eximer-forming lipid (Lopez *et al*., 2017), to measure the fluidity of WT and Δ*esxC* membranes. Compared to the WT, the cytosolic membrane of the Δ*esxC* mutant showed a mild but statistically significant increase in rigidity (Fig. 1C). Furthermore, we stained WT and with a fluorescent membrane dye DiI12C, which has been used detect regions of increased fluidity in bacteria (Saeloh *et al*., 2018). As reported previously for *S. aureus*, we observe few fluid regions in the WT, and an uneven staining of the membrane. In contrast, a very weak staining of the membranes was observed for the Δ*esxC* mutant, (Fig 1 D and E), which may indicate a decrease in membrane fluidity.

Overall, the data indicate that EsxC contributes to *S. aureus* cell surface structure and membrane fluidity.

### *S. aureus esxC* and *essC* mutants are more sensitive to antimicrobial fatty acids

Multiple studies have reported the activation of the T7SS, when *S. aureus* is grown in presence of host FAs (Ishii *et al*., 2014, Kenny *et al*., 2009, Lopez *et al*., 2017). Given our findings that the *esxC* mutant has membrane defects and that the *esxC* deletion affects the surface abundance of IsdA, which is essential for *S. aureus* resistance to antimicrobial fatty acids (Clarke *et al*., 2007), we cultured WT USA300 and its isogenic *essC* or *esxC* mutant in the presence of an unsaturated FA (C18:2), linoleic acid (LA), at a concentration (80 µM) that still allows WT growth. Bacteria were also cultured in parallel in presence of stearic acid (SA), a saturated C18:0 FA. The T7SS mutants displayed impaired growth in presence of LA but not SA, as measured by optical density (OD) (Fig. 2A) or colony forming units (CFU) (Fig. 2B). Importantly, Δ*esxC* complemented with a plasmid containing the *esxC* gene reverted to the WT phenotype (Fig. 2C). The increased susceptibility of T7SS mutants to antimicrobial fatty acids was not restricted to linoleic acid; when cultured in the presence of arachidonic acid, another unsaturated FA (C20:4), growth of Δ*esxC* and Δ*essC* were inhibited more as compared to the WT (Fig. S4)

**Fig. 2.**
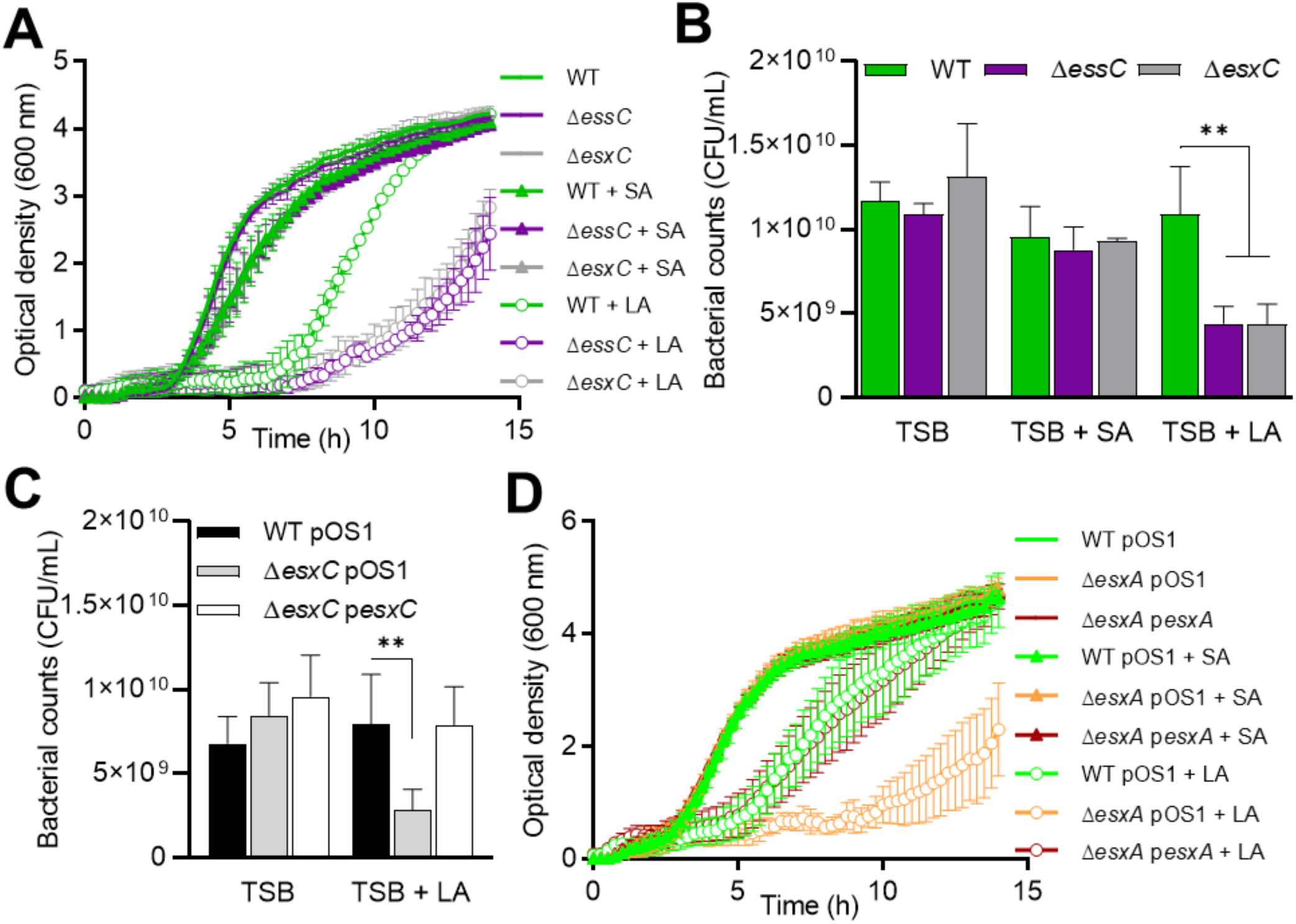
Enhanced *S. aureus* growth inhibition by linoleic acid in T7SS mutants. **A.** *S. aureus* WT USA300, Δ*essC*, and Δ*esxC* were grown in TSB or TSB supplemented with 80 µM of either linoleic (LA) or stearic acid (SA). Means ± standard error of the mean (SEM) are shown. **B.** After 14h growth as described in (A) bacteria were serially diluted, and CFU were determined. Mean values are presented, and the error bars represent SD. *n* = 3, ** indicates *P* < 0.01 using one-way ANOVA with Dunnett’s test. **C.** USA300 WT with the empty pOS1 plasmid (WT pOS1) and USA300 JE2 *esxC* mutant with either pOS1 (Δ*esxC* pOS1) or pOS1-*esxC* (Δ*esxC* pOS1-*esxC*) were grown in TSB or TSB + 80 µM LA as described in (A) followed by CFU estimation. Mean values are shown; error bars represent SD. *n* = 5, ** indicates *P* < 0.01 using one-way ANOVA with Dunnett’s test. **D.** Newman WT with the empty pOS1 plasmid (WT pOS1) and Newman *esxA* mutant with either pOS1 (Δ*esxA* pOS1) or pOS1-*esxA* (Δ*esxA* p*esxA*) were grown in TSB or TSB + 40 µM LA or SA. Means ± standard error of the mean (SEM) are shown, *n* = 4.

### T7SS substrates contribute to *S. aureus* resistance to LA toxicity

Next, we investigated whether T7SS proteins other than *essC* and *esxC* contributed to *S. aureus* growth in presence of LA. Mutants lacking two other substrates, Δ*esxA* and Δ*esxB*, were grown in presence of FAs. Both mutants grew slower than the WT USA300 (Fig. S5A). To ensure that the increased sensitivity observed for the T7SS mutants was not strain specific, RN6390 Δ*essC* or Δ*esxC* and Newman Δ*esxA* or Δ*esxB* mutants were tested. Similar to the USA300 mutants, the growth of all these T7SS mutants was also impacted in the presence of LA (Fig. S5B and C). The growth defect in Newman Δ*esxA* was abrogated upon complementation (Fig. 2D). Of note, the Newman WT was readily inhibited by a lower concentration of LA (40 µM), which is in agreement with the lower T7SS expression levels in this strain compared to USA300 (Anderson *et al*., 2013, Kneuper *et al*., 2014). We conclude that a functional T7SS plays a role in *S. aureus* resistance to LA toxicity.

### T7SS is required for maintaining membrane integrity in presence of LA

To study the mechanisms involved in T7SS mediated protection to LA toxicity, further studies were performed using strains lacking esxC, a representative T7SS effector, or essC the main T7SS transporter. As our membrane and surface proteome analyses with Δ*esxC* suggested that the bacterial envelope is altered upon T7SS defect, we wanted to test if LA-mediated growth inhibition was due to an increased binding of LA to T7SS mutants. To do this, we chemically engineered LA to produce an azide functionalised LA (*N^6^*-diazo-*N^2^*-((9Z,12Z)-octadeca-9,12-dienoyl)lysine, N_3_-LA) or azide-LA (Fig. 3A). After incubating bacteria with azide-LA, click-chemistry with an alkyne dye (Click-iT™ Alexa Fluor™ 488 sDIBO alkyne) was used to stain azide-LA associated with bacteria. There were no obvious differences in the fluorescence from Δ*essC* and Δ*esxC* compared to the WT (Fig. 3B), suggesting that T7SS components are not involved in binding or sequestering LA. However, when bacteria treated with azide-LA were stained with propidium iodide (PI), a good indicator of membrane integrity, Δ*essC* and Δ*esxC* displayed a more intense PI staining compared to the WT (Fig. 3C and D). Similarly, when WT and mutants were stained with PI and SYTO 9 after treatment with 80 µM unlabelled LA, we observed an increased PI staining (Fig. 4A and B) and therefore lower SYTO 9 / PI (Live/Dead) ratio for both mutants (Figure 4C). These data suggest that an intact T7SS helps *S. aureus* to maintain its membrane integrity when faced with the detergent-like effects of unsaturated FAs.

**Fig. 3.**
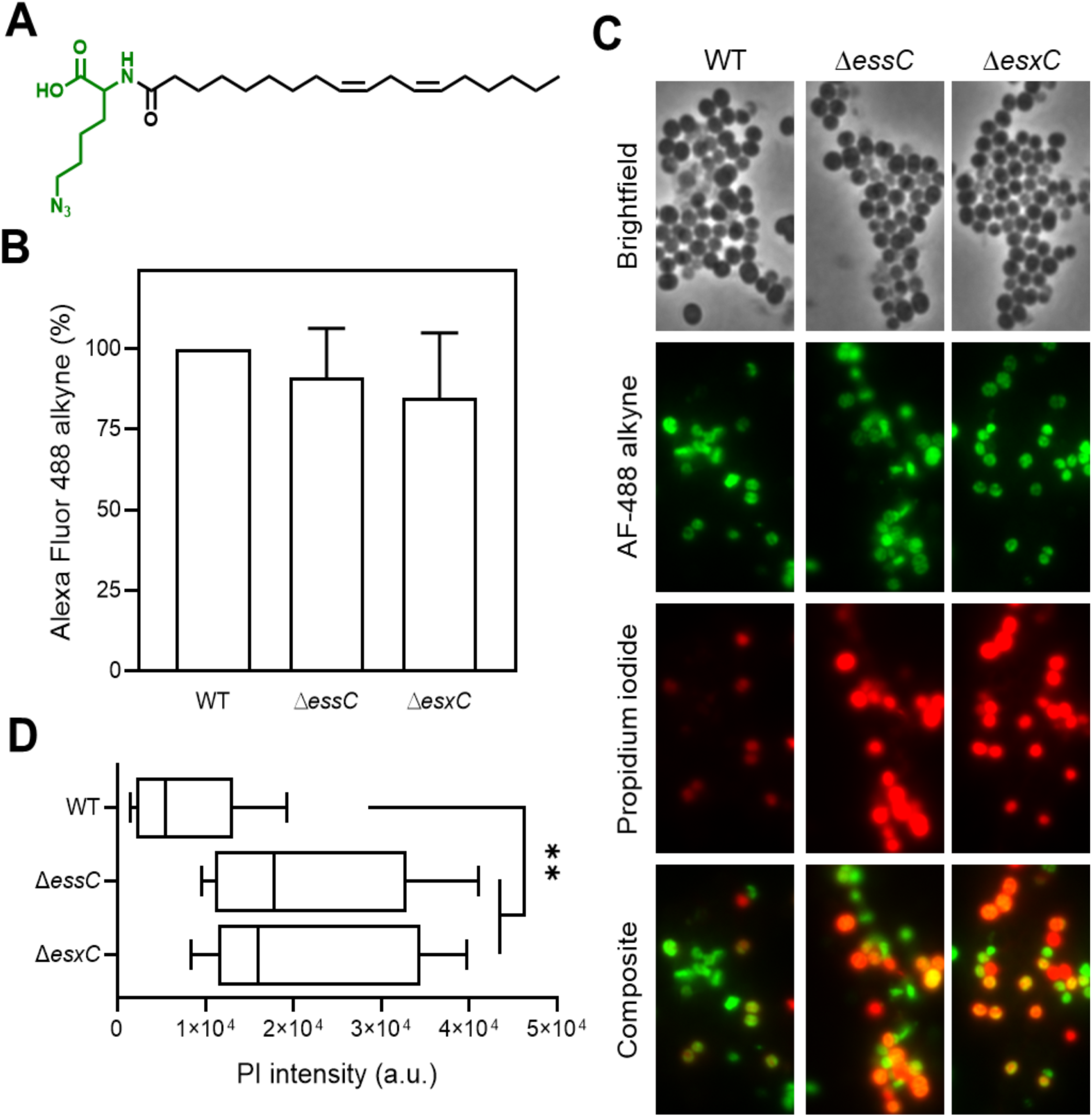
T7SS mutants display increased membrane permeability upon LA binding. **A.** Chemical structure of azide functionalised linoleic acid (azide-LA; *N^6^*-diazo-*N^2^*-((9Z,12Z)-octadeca-9,12-dienoyl)lysine, N_3_-LA). Highlighted in green is the azido lysine. **B.** *S. aureus* USA300 WT, Δ*essC*, and Δ*esxC* were grown with shaking in TSB to OD_600_ of 1.0. Bacteria were then stained for 15 min with 10 µM azide-LA prior to labelling for 1 h with alkyne Alexa Fluor 488. Mean percentage of fluorescence values relative to WT (100%) are presented; error bars represent SD, *n* = 5. **C.** Micrographs of bacteria grown in TSB and treated as described in (B) and additionally stained with propidium iodide (PI). **D.** ImageJ was used to quantitate PI fluorescence of bacterial clusters from 12 different fields per strain. Each box-and-whisker plot depicts the minimal and maximal PI intensities, the median is the vertical bar inside the box, which is delimited by the lower and upper quartiles. ** indicates *P* < 0.01 using one-way ANOVA with Dunnett’s test.

**Fig. 4.**
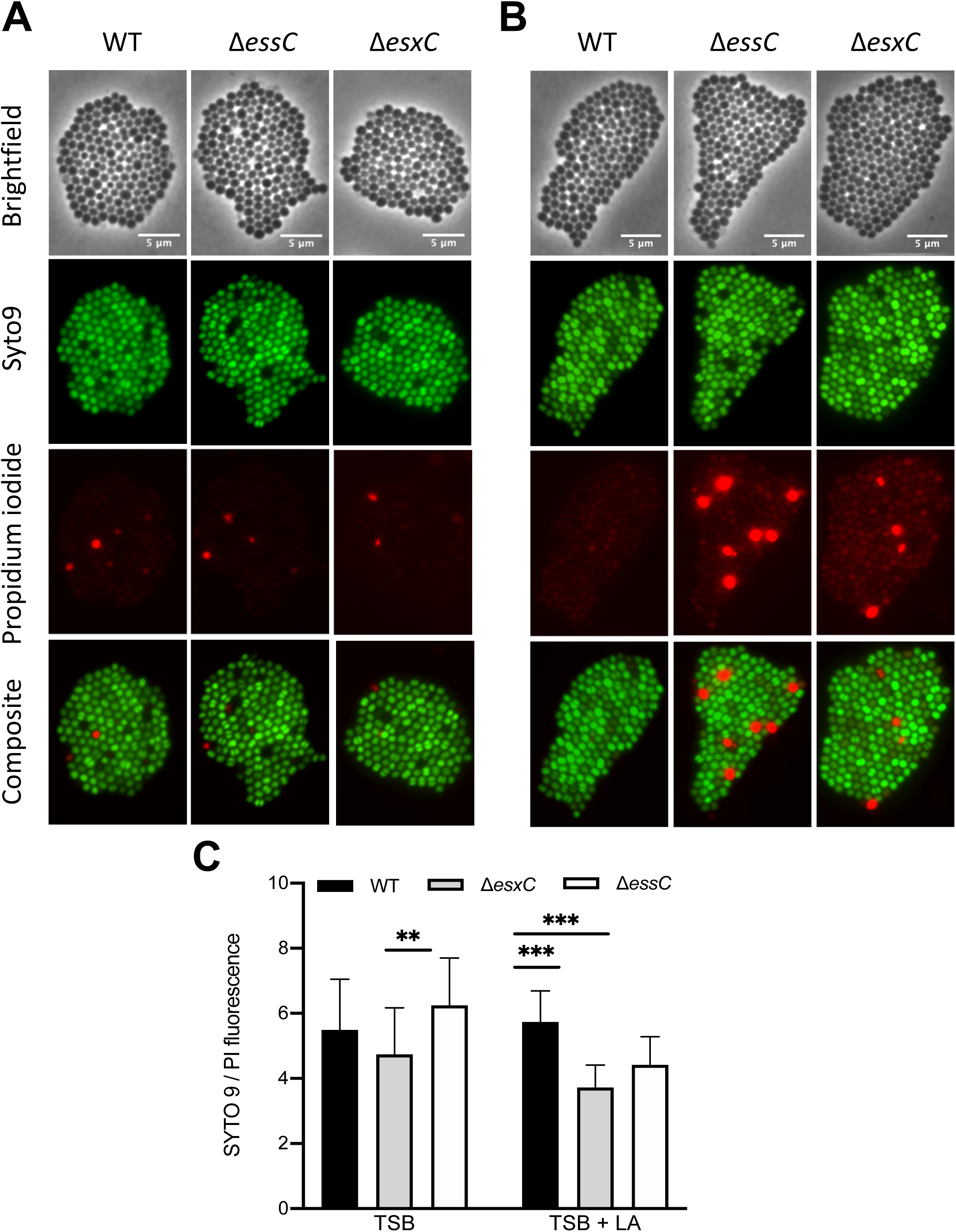
T7SS mutants display increased PI staining when treated with LA. Live/Dead staining of *S. aureus* USA300 WT, Δ*essC* or Δ*esxC* mutants after growth to OD_600_ of 1.0, without (**A**) or with treatment with 80µM (**B**) linoleic acid. Images are representative of 3 independent experiments. **C**. The ratio of SYTO 9: PI fluorescence (live:dead cells) of 25 different fields per strain was quantitated with ImageJ. Means ± SD are shown, *n* = 3; *** indicates *P <* 0.001, ** indicates *P <* 0.01 using a one-way ANOVA with Tukey’s multiple-comparison test

### LA-incorporation into membrane phospholipids is modulated by T7SS

As the T7SS mutants have more compromised cell membranes in presence of LA, we next investigated if membrane lipids were altered in the T7SS mutants. Lipids from WT USA300 and T7SS mutants were analysed by high-performance liquid chromatography (HPLC)-mass spectrometry (MS) in negative ionisation mode. As reported previously (Delekta *et al*., 2018, Parsons *et al*., 2011), phosphatidylglycerol (PG) was the major phospholipid present in the membrane of WT grown in TSB (Fig. S6A). Δ*essC* and Δ*esxC* grown with or without 10 µM LA [(a concentration that has been previously shown to be sub-inhibitory for USA300 (Lopez *et al*., 2017)] displayed lipid profiles similar to that of WT (Fig. S6A and B). Notably, PG molecular species were significantly altered upon growth in LA-supplemented TSB for WT (Fig. 5A), Δ*essC* (Fig. S6C) and Δ*esxC* (Fig. 5B). Three new LA-specific PG species with mass to charge ratios (m/z) 731 (C33:2), 759 (C35:2), and 787 (C37:2) appeared to contain LA (C18:2) or its elongated C20:2 or C22:2 versions, as revealed by their fragmentations (Fig. S7A, B and C). PG species containing exogenous, unsaturated FAs were also present in Δ*essC* and Δ*esxC*. However, LA (C18:2)-containing PG species (C33:2) were less abundant in the *esxC* mutant compared to WT (Fig. 5C). A similar trend, although statistically non-significant (*P* > 0.05), was observed for C20:2- and C22:2-containing PG species (Fig. S7D and E), and when all the unsaturated exogenous PG species were combined (Fig. 5D). The data suggest that a T7SS defect may compromise the incorporation and elongation of LA in *S. aureus* membranes.

**Fig. 5.**
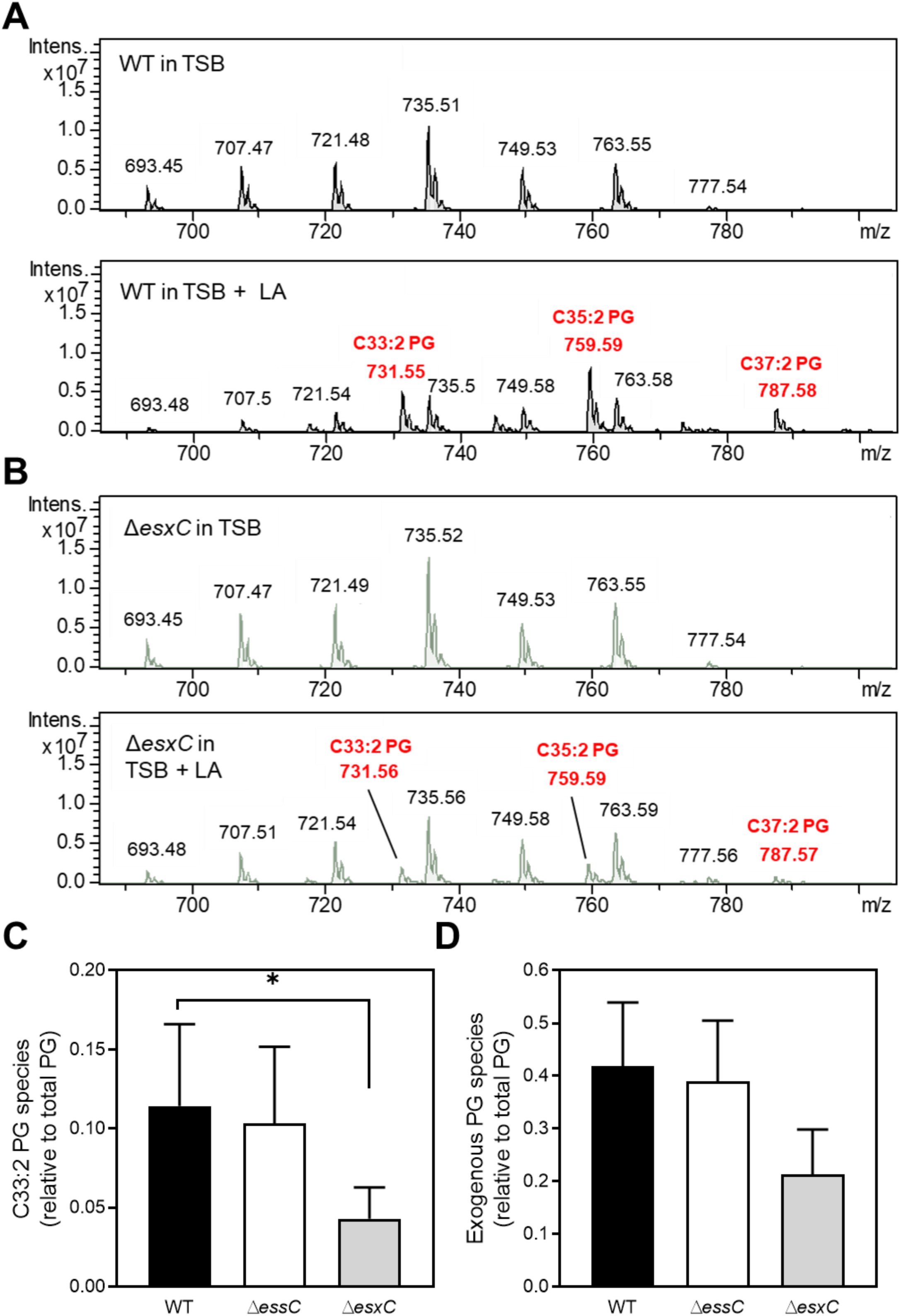
T7SS mutants are less able to incorporate LA into their phospholipids. Representative HPLC chromatograms of native phosphatidylglycerol (PG) species of *S. aureus* USA300 JE2 WT (**A**) or Δ*esxC* (**B**) grown in TSB (top panel) or in TSB supplemented with 10 µM LA (bottom panel), in negative ionisation mode. Relative quantification of the indicated PG species containing an unsaturated FA in WT, Δ*essC* and Δ*esxC*. C18:2-containing PG species (**C**) and total unsaturated exogenous PG species (**D**) are presented as ratios of total PG species. Mean values are shown; error bars represent SD. *n* = 3, * indicates *P* < 0.05 using one-way ANOVA with Dunnett’s test.

### T7SS mutations affect the total cellular content and *S. aureus* responses to LA

In order to gain further insight into T7SS-mediated modulation of proteins involved in FA incorporation and membrane homeostasis in presence of LA, we used an unbiased proteomic approach to study protein profiles of WT USA300, Δ*essC*, and Δ*esxC* grown exponentially with or without 10 µM LA. Of note, WT and both these T7SS mutants grew similarly in presence of up to 40 µM LA (Fig. S8).

#### WT vs T7SS mutants in absence of LA treatment

Interestingly, Δ*essC* or Δ*esxC* cultured in TSB readily displayed proteins with changed abundance when compared to the WT, with 37 and 24 proteins significantly (*P* < 0.05) altered in Δ*essC* and Δ*esxC*, respectively. Similarly, 14 proteins were differentially abundant in both Δ*essC* and Δ*esxC* (Fig. 6A and B). These included proteins associated with signal transduction (LytR and ArlR), the CW (acetyltransferase GNAT, FnbB and MazF), DNA repair (MutL and RadA), nucleotide binding (ATP-grasp domain protein and YqeH), hydrolysis (amidohydrolase), cell stress response [universal stress protein (Usp) family], or were uncharacterised (A0A0H2XGJ8, YbbR and lipoprotein) (Fig. 6B and Table 1). Of the 33 proteins changed only in Δ*essC* (23 proteins) or Δ*esxC* (10 proteins), nearly 40% (13 proteins) were associated with oxidation-reduction and other metabolic processes. Ten membrane proteins were more abundant in Δ*essC* (Table 1), which included SrrB, a membrane protein that is activated by impaired respiration (Mashruwala *et al*., 2017), and whose gene expression increased 6 times upon growth in presence of LA (Lopez *et al*., 2017). SrrB was also detected at higher levels in the *esxC* mutant although the increase was non-significant (*P* = 0.07).

**Fig. 6.**
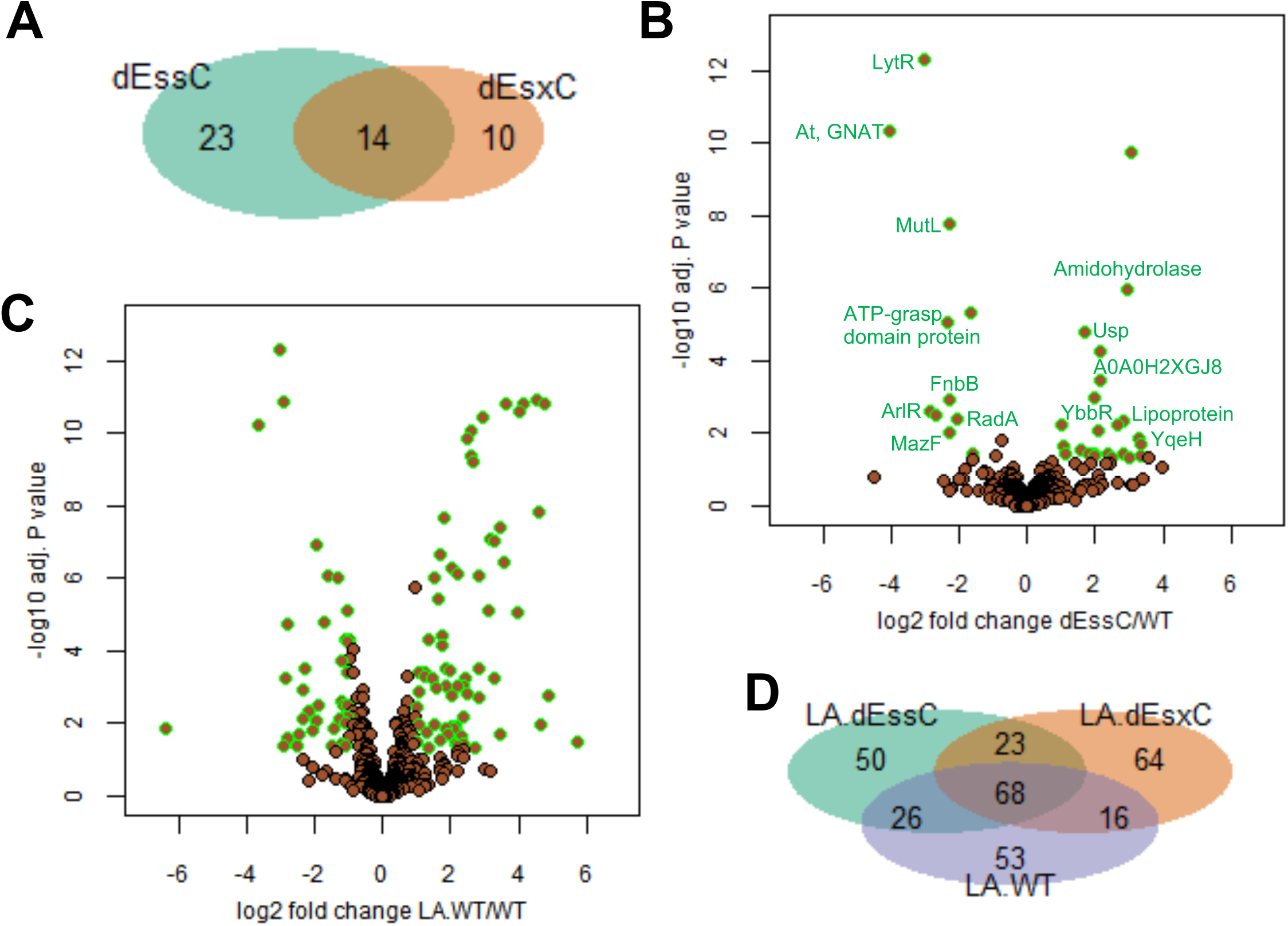
Quantitative proteomics shows altered cellular content and bacterial response to LA in T7SS mutants. *S. aureus* USA300 WT and mutants (Δ*essC* and Δ*esxC*) were grown in TSB or TSB supplemented with LA. **A.** Venn diagram showing the number of proteins with altered abundance compared to WT specific to Δ*essC* (23) or Δ*esxC* (10), and common to Δ*essC* and Δ*esxC* (14). **B.** The fourteen proteins that are similarly changed in Δ*essC* and Δ*esxC* mutants are highlighted on a volcano plot. **C.** Volcano plot showing the extensive change in the LA-treated WT compared to WT. **D.** Venn diagram displaying the numbers of proteins with altered relative abundance upon LA challenge of WT (LA.WT), Δ*essC* (LA.dEssC) or Δ*esxC* (LA.dEsxC) compared to the respective untreated samples.

**Table 1.**
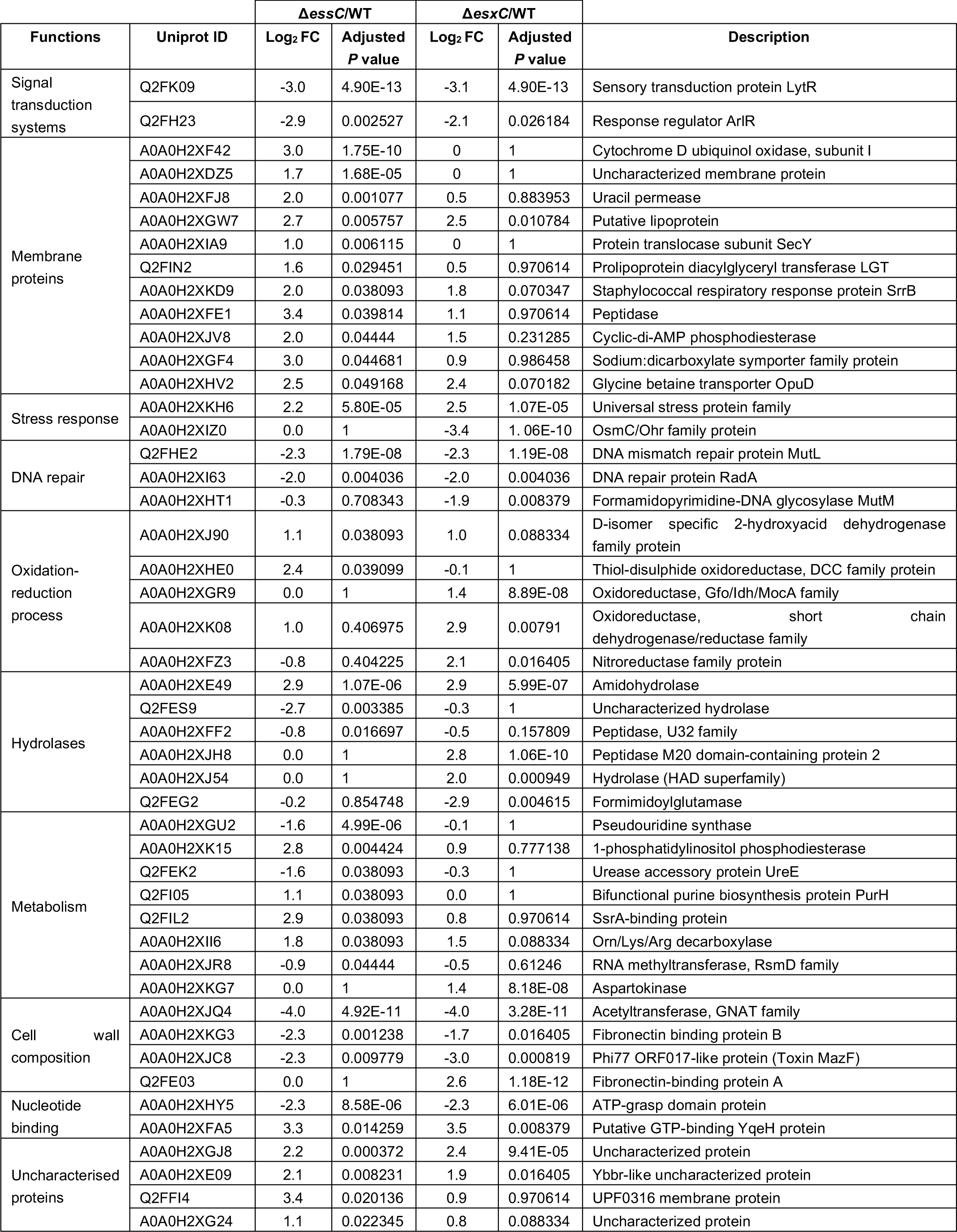
Proteins with changed abundance in Δ*essC* and Δ*esxC* mutants relative to the WT USA300 JE2.

#### WT vs T7SS mutants in presence of LA

We then compared the proteomic profiles of LA-treated strains (WT, Δ*essC* or Δ*esxC*) with their untreated counterparts. Clearly, the principal component analysis revealed that the differences due to the genetic makeup (WT or T7SS mutants) were less prominent than the dramatic changes induced by LA (Fig. S9). These changes are exemplified for the WT; 163/1132 proteins identified had an altered relative abundance upon growth with LA (Fig. 6C). 167 and 171 proteins were changed (*P* < 0.05) in Δ*essC* and Δ*esxC*, respectively, in response to LA, of which ∼ 40% (68 proteins) were common to these mutants and their WT (Fig. 6D). At least 30% of the significantly changed proteins (*P* < 0.05) were unique to WT (53 proteins), Δ*essC* (50 proteins), or Δ*esxC* (64 proteins) (Fig. 6D), suggesting that each strain responds differently to LA. However, almost all proteins (13/14 proteins) that were similarly deregulated in Δ*essC* and Δ*esxC* grown without LA (Fig. 6B) were modulated in presence of LA (highlighted in bold in Dataset S1). Proteins that were less abundant in both mutants were, upon LA treatment, either increased to WT levels (MutL, acetyltransferase GNAT, Toxin MazF, and ATP-grasp domain protein), or were unchanged in the mutants and decreased in the LA-treated WT (LytR and FnbB) (Dataset S1). Likewise, proteins with increased amounts in Δ*essC* or Δ*esxC* were: (i) downregulated to WT levels in response to LA (putative lipoprotein A0A0H2XGW7), (ii) unaltered in both mutants and upregulated in WT (Usp, amidohydrolase, and YbbR), (iii) or further increased in the *essC* mutant and strongly upregulated in WT (A0A0H2XGJ8) (Dataset S1). In sum, except for ArlR and RadA that were conversely regulated in all strains after LA treatment, proteins similarly deregulated in Δ*esxC* and Δ*essC* were returned to similar levels in response to LA. A similar trend was observed for 15/23 and all 10 proteins exclusively more or less abundant in Δ*essC* and Δ*esxC*, respectively.

#### Altered molecular functions in presence of LA

We then used QuickGO (a web-based tool for Gene Ontology searching) (Binns *et al*., 2009) to retrieve GO terms associated with the ten most significantly upregulated proteins in LA-treated WT (Dataset S1). Strikingly, 9/10 proteins had a hydrolase or an oxidoreductase activity. A comprehensive, statistical analysis showed a clear enrichment of 8 specific molecular functions (*P* < 0.05) in at least one strain (WT or T7SS mutants) (Fig. 7A). Oxidoreductase and hydrolase activities were enhanced in LA-treated WT, while Δ*essC* and Δ*esxC* were less able to upregulate proteins with these molecular functions. Flavin adenine dinucleotide (FAD)-binding, which plays a role in oxidation-reduction and FA metabolic processes, was similarly more enriched in the LA-treated WT. In contrast, transferase activity, which is linked to CW synthesis, was induced more in T7SS mutants compared to the WT. Molecular functions that are decreased upon LA challenge were also determined (Fig. 7B). In agreement with reduced intracellular ATP levels following membrane damage by antimicrobial FAs (Cartron *et al*., 2014), genes with the ATP-binding function (mainly ATP-binding ABC transporters) were strongly inhibited in the WT. ATP-dependent lyases were also repressed in the WT. On the contrary, T7SS mutants were less able to modulate ATP-binding proteins. Instead, a strong inhibition of ribosomal constituents and other translation-related components was seen (Fig. 7B).

**Fig. 7.**
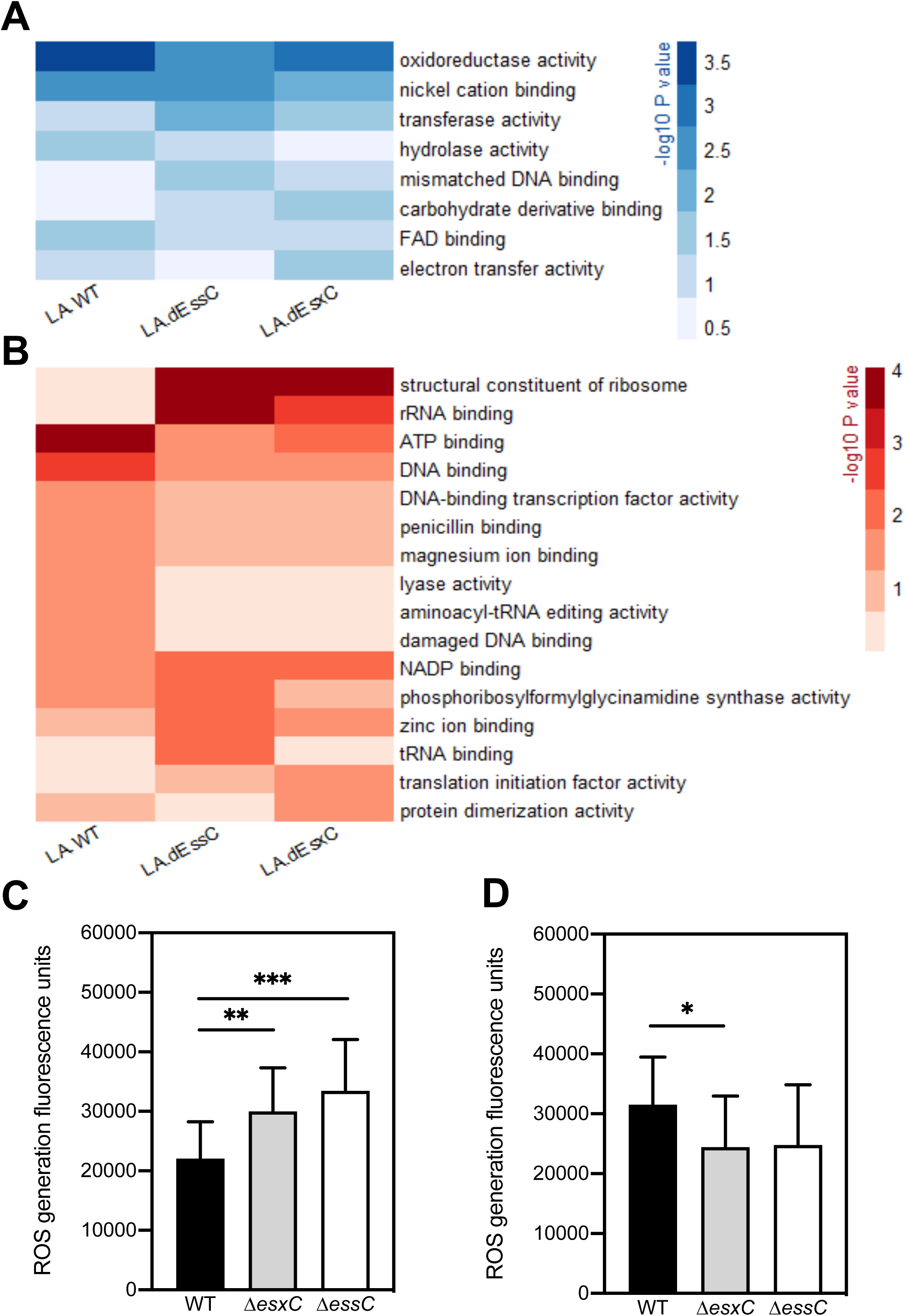
An altered oxidoreductive response in T7SS mutants in response to LA. Heatmaps depicting the *P* values of enriched (**A**) or diminished (**B**) molecular functions following a gene set analysis based on GO (gene ontology) annotations. Molecular functions that are changed in at least one strain (*P* < 0.05) following growth in presence of LA are shown. The shades of blue (**A**) or red (**B**) correspond to – log_10_ (*P* value). ROS levels were measured in cultures of *S. aureus* USA300 JE2 WT, Δ*essC* or Δ*esxC* grown to OD_600_ of 1.0 treated (**C)** with or without LA (**D**) using DCF reagent. Means ± SD are shown N=5. * indicates *P <* 0.05, ** indicates *P <* 0.01, *** indicates *P <* 0.001 using the Kruskal-Wallis rank test.

To test the oxidoreductive states of the WT and the mutants, we stained bacteria with dichlorofluorescin (DCF), which detects reactive oxygen species (George *et al*., 2019). Reflecting the changes seen in the proteomics data, when treated with 10 µM LA there is an increase in the ROS generated in the T7SS mutants compared to the WT (Fig 7C). However, in bacteria grown without LA, the mutants have slightly less or no change in the ROS generated compared to WT (Figure 7D). Taken together, our proteomic analyses reveal that the lack of T7SS induces altered membrane and metabolic states indicative of oxidative stress responses. While the WT shows a multifaceted response to mitigate LA-induced damage on the bacterial membrane, such responses are clearly altered in the absence of the T7SS.

## Discussion

Host fatty acids (FAs) play a crucial role in the host defence to *S. aureus* infections. *S. aureus* is particularly sensitive to unsaturated FAs, which are abundant in the human skin (Clarke *et al*., 2007, Parsons *et al*., 2012, Arsic *et al*., 2012, Kelsey *et al*., 2006, Kenny *et al*., 2009). We report here that the T7SS, an important component of *S. aureus* virulence arsenal, is critical in modulating the response to antimicrobial host FAs by maintaining the bacterial cell membrane integrity. Specifically, we demonstrate that a T7SS substrate, EsxC, impacts *S. aureus* membrane properties. A functional T7SS enables bacteria to mitigate LA-induced toxicity and grow better than mutants with a compromised T7SS. In the absence of T7SS components, LA is less incorporated into membrane phospholipids and enhances cell membrane damage, while bacteria are unable to activate adaptive mechanisms involved in LA resistance, as indicated by cellular proteomics.

Although several studies have shown multiple interactions between the staphylococcal T7SS components, the precise molecular architecture of this system remains unclear. EsxC (previously EsaC) was first described as a secreted protein (Burts *et al*., 2008). However, in keeping with prior studies, we found that EsxC can also localize within staphylococcal membranes (Bobrovskyy *et al*., 2018, Kneuper *et al*., 2014). Based on available data, EsxC is likely to be associated to EsxA, EsaD, or EsaE on the membrane (Anderson *et al*., 2013, Anderson *et al*., 2017, Cao *et al*., 2016). Additionally, the reduced EsaA protein levels that we found on the surface of *S. aureus* USA300 Δ*esxC* combined with prior observations of diminished EsxC protein levels in RN6390 Δ*esaA* membranes (Kneuper *et al*., 2014) suggest that EsxC may interact with EsaA, a key component of the T7SS core (Aly *et al*., 2017), in *S. aureus* membranes.

T7SS integral membrane proteins interact with the flotillin homolog FloA within functional membrane microdomains (FMMs) (Mielich-Suss *et al*., 2017). FMMs were recently shown to contain staphyloxanthin derivatives (Garcia-Fernandez *et al*., 2017), which are known to increase membrane rigidity (Chamberlain *et al*., 1991, Sen *et al*., 2016, Tiwari *et al*., 2018). Hence, well-structured FMMs may play a role in *S. aureus* membrane fluidity as reported for *Bacillus subtilis* (Bach & Bramkamp, 2013). Interestingly, mutations or treatments affecting *S. aureus* membrane fluidity also modulate T7SS transcription (Lopez *et al*., 2017, Ishii *et al*., 2014, Parsons *et al*., 2014). Hence, there appears to be a link between membrane fluidity and T7SS in *S. aureus*. Also, the state of the membrane was suggested to trigger the production of T7SS (Lopez *et al*., 2017). Remarkably, deletion of *esxC* led to a mild increase in bacterial membrane rigidity and membrane defects (Fig. 1), suggesting membrane modulation by the T7SS. Possibly, interactions between FloA and T7SS are perturbed upon *esxC* deletion and affect *S. aureus* FMMs. We surmise that a functional T7SS helps *S. aureus* to maintain its membrane architecture.

The current cellular proteomics data reveal that the abundance of more proteins is altered in Δ*essC* (37) than *esxC* (24) in comparison to *S. aureus* WT, which is in keeping with the greater importance of EssC as the conserved driving force of the T7SS (Warne *et al*., 2016). Importantly, almost 60% of proteins deregulated in Δ*esxC* are similarly affected in Δ*essC*, strongly suggesting that any modification of the T7SS core leads to a similar staphylococcal response. Surprisingly, proteins with altered abundance in USA300 Δ*essC* were distinct to the ferric uptake regulator (Fur)-controlled genes differentially expressed in RN6390 Δ*essC* (Casabona *et al*., 2017b). This discrepancy might be due to strain differences, including *rsbU* defect in RN6390 that impairs SigB activity (Cassat *et al*., 2006, Giachino *et al*., 2001). Nevertheless, given the known role of Fur in oxidative stress resistance (Horsburgh *et al*., 2001, Johnson *et al*., 2011), both mutants may display an altered oxidative status following *essC* deletion. *S aureus* RN6390 also differentially expresses redox-sensitive genes in absence of EsaB (Casabona *et al*., 2017a). Also, since the T7SS substrate EsxA is upregulated in response to hydrogen pyroxide (Casabona *et al*., 2017b), one could speculate that lack of T7SS stimulates an oxidative stress response. A further indication of altered physiological states of Δ*essC* and Δ*esxC* was the decreased abundance of the two-component regulatory system proteins, LytSR, ArlSR and SrrAB, which was consistent with down-regulation of *lytR* transcription observed previously in the absence of *arlR* (Liang *et al*., 2005). Importantly, the *S. aureus* response to antimicrobial FAs includes downregulation of *lytRS* (Kenny *et al*., 2009, Neumann *et al*., 2015), and upregulation of *srrB* (Lopez *et al*., 2017). Given that LytSR is involved in bacterial surface and membrane potential modulation (Patton *et al*., 2006, Groicher *et al*., 2000), T7SS defects are likely to result in an altered cell envelope.

It is striking that the staphylococcal T7SS is strongly upregulated in presence of sub-inhibitory concentrations of LA (Kenny *et al*., 2009, Lopez *et al*., 2017). FAs with more cis double bonds, which are more toxic toward *S. aureus* (Parsons *et al*., 2012), are also more potent T7SS activators (Lopez *et al*., 2017). Our current study interestingly suggests a protective role of T7SS against LA toxicity. Previously described *S. aureus* antimicrobial FA (AFA) resistance mechanisms, including IsdA or wall teichoic acid-mediated modulation of cellular hydrophobicity (Clarke *et al*., 2007, Kohler *et al*., 2009, Parsons *et al*., 2012, Moran *et al*., 2017), and AFA detoxification with the efflux pumps Tet38 and FarE (Alnaseri *et al*., 2015, Truong-Bolduc *et al*., 2014), do not appear to explain the increased susceptibility of T7SS mutants to LA, as indicated by cellular proteomics. In line with a role for T7SS in the oxidative stress response, T7SS mutants were less able to prime their redox-active proteins in response to LA-induced oxidative stress. Instead, to cope with LA, they appear to rely on strong inhibition of the protein synthesis machinery, which is reminiscent of the stringent response (Geiger *et al*., 2012).

The Fak pathway which is required for incorporation of exogenous FA into membrane phospholipids via a two-component fatty acid kinase (Fak) (Nguyen *et al*., 2016, Parsons *et al*., 2012, Parsons *et al*., 2014), was reported to be important for T7SS activation by unsaturated FA (Lopez *et al*., 2017). FakB1 and FakB2, bind to FAs, and FakB-bound FAs are phosphorylated by FakA prior to their incorporation (Parsons *et al*., 2014). In presence of inhibitory concentrations of unsaturated FA like LA, while FA is incorporated into the membrane lipids, free LA causes pore formation that leads to bacterial lysis (Greenway & Dyke, 1979). Our lipidomic analyses revealed that in the absence of T7SS, bacteria were less able to incorporate LA into their phospholipids (Figure 5), and displayed an increased membrane permeability in presence of LA. However, it seems counterintuitive that LA incorporation was impacted more in Δ*esxC* than in Δ*essC* given the central role of EssC in T7 secretion (Burts *et al*., 2005, Jager *et al*., 2018, Zoltner *et al*., 2016). It is possible that EsxC that accumulates in the membrane (Fig. 1A) in the absence of protein secretion by the main transporter, EssC, mediates partial LA incorporation in the *essC* mutant; secretion per se may not be required for LA incorporation. It is also worth noting that transcript levels of *esxC*, and not *essC*, were strongly upregulated in a *S. aureus fakA* mutant (Parsons *et al*., 2014). As protein levels of Fak proteins in the T7SS mutants stay unaltered in presence or absence of LA, suggesting no T7SS-mediated regulatory control of the Fak pathway, we speculate that EsxC and other interdependent T7SS substrates play an important role in facilitating Fak function in *S. aureus* membranes, either by mediating recruitment or targeting of Fak proteins to the membrane. Our findings warrant further investigations into molecular mechanisms underlying T7SS-mediated FA incorporation within staphylococcal membranes.

The increased susceptibility of T7SS mutants to LA might explain why they are less virulent in environments rich in LA and other AFAs like the mouse lungs (Δ*essC*) (Ishii *et al*., 2014), abscesses (Δ*esxC* and Δ*esaB*), liver and skin (Δ*essB*) (Wang *et al*., 2016, Lopez *et al*., 2017). Previous research showing T7SS induction by host-derived FAs further supports the importance of T7SS in such environments (Lopez *et al*., 2017, Ishii *et al*., 2014). Taken together, we conclude that T7SS plays a key role in modulating the *S. aureus* cell membrane in response to toxic host FAs. Although, at present, it is unclear how T7SS contributes to staphylococcal membrane architecture, T7SS interaction with the flotillin homolog FloA within functional membrane microdomains (Mielich-Suss *et al*., 2017) corroborates the idea that T7SS proteins interact with many other proteins to modulate *S. aureus* membranes. Indeed, our data also suggest that blocking T7SS activity would make *S. aureus* more vulnerable to AFAs, a key anti-staphylococcal host defence, thus making T7SS a very attractive drug target.

## Experimental procedures

### Bacterial strains and growth conditions

*S. aureus* strains used are listed in Table S3, and were grown aerobically in tryptic soy broth (TSB) overnight (O/N) at 37°C for each experiment unless stated otherwise. For complemented *S. aureus* strains, TSB was supplemented with 10 µg/mL chloramphenicol.

### Construction of bacterial mutants

The primers used are listed in Table S4. In-frame deletion of *essC* or *esxC* was performed as described previously (Bae & Schneewind, 2006). Briefly, 1-kb DNA fragments up and downstream of the targeted gene sequence were PCR-amplified from USA300 LAC JE2 chromosomal DNA, and both PCR products fused via SOEing (splicing by overlap extension)-PCR. The 2-kb DNA fragment obtained was cloned into pKORI, and used for in-frame deletion. Putative mutants were screened by PCR-amplification of a fragment including the gene of interest, whose deletion was confirmed by Sanger sequencing. Further, to confirm that successful mutants did not have any additional mutations, lllumina whole genome sequencing was performed on libraries prepared with the Nextera XT kit and an Illumina MiSeq® instrument following manufacturers’ recommendations. For complementation, full-length *esxC* gene was cloned onto pOS1CK described previously (Korea *et al*., 2014).

### Membrane fluidity assay

O/N bacterial cultures were diluted to an OD_600_ of 0.15 in TSB, and were grown to an OD_600_ of 1 (OD1). Bacteria were washed with PBS prior to treatment for 30 min at 37°C with 37.5 µg/mL lysostaphin in PBS containing 20% sucrose. The spheroblasts were then centrifuged at 8000 × *g* for 10 min, and the pellet resuspended in the labelling solution (PBS, 20% sucrose, 0.01% F-127, 5 µM pyrene decanoic acid). The incubation in the dark was done for 1h at 25°C under gentle rotation. PBS supplemented with 20% sucrose was used to wash the stained spheroblasts that were afterwards transferred to 96-well plates for fluorescence measurements as previously described (Lopez *et al*., 2017).

### Triton X-100 lysis assays

Whole cell autolysis assays were performed as described elsewhere with a few modifications (Mashruwala *et al*., 2017). Specifically, OD1-grown *S. aureus* USA300 JE2 WT, *essC* and *esxC* mutants were extensively washed with PBS followed by ice-cold water, and resuspended in PBS with 0.1% Triton X-100 to an OD_600_ of 0.7. Subsequently, the samples were incubated with shaking at 37°C for 2h, after which bacteria were diluted with PBS and plated for CFU determination.

### FM1-43 staining

*S. aureus* WT and mutant strains grown to OD_600_ of 1.0 were centrifuged, and pellets were resuspended in PBS supplemented with 1 μg/mL FM1-43 (Invitrogen). After incubation for 15 min in the dark at 37°C with shaking, cells were washed once with PBS. Fluorescence was quantified using a FLUOstar OMEGA plate reader (BMG Labtech, UK) at excitation and emission wavelengths of 482 nm and 620 nm, respectively.

### DiIC12 staining

O/N bacterial cultures were diluted to an OD_600_ of 0.15 in TSB with 1 µg/mL 1,1′-didodecyl-3,3,3′,3′-tetramethylindocarbocyanine perchlorate (DiIC12) (Invitrogen). Cultures were grown to an OD_600_ of 1.0, centrifuged and washed twice with fresh TSB. Samples were spotted on to agarose pads and imaged using a Leica DMi8 widefield microscope (Leica Microsystems, UK). Acquired images were analysed with the ImageJ processing package, Fiji.

### Growth curves

O/N bacterial cultures were diluted to an OD_600_ of 0.05 in plain TSB or TSB supplemented with fatty acids. Bacteria were then grown in a 96-well plate with shaking, and the OD_600_ was measured every 15 mins with a FLUOstar OMEGA plate reader (BMG Labtech, UK).

### Synthesis of azide functionalized linoleic acid

A 2-step synthesis was used to obtain *N^6^*-diazo-*N^2^*-((9Z,12Z)-octadeca-9,12-dienoyl)lysine, N_3_-LA (azide-LA). LA was first functionalized with *N*-hydroxysuccinimide (NHS) in anhydrous dimethyl formamide (DMF) in presence of *N*-(3-dimethylaminopropyl)-*N*’-ethylcarbodiimide hydrochloride. The solvent was then removed and replaced by dicholoromethane (DCM), following which the reaction mixture was washed with water and dried over magnesium sulphate. The product, 2,5-dioxopyrrolidin-1-yl (9Z,12Z)-octadeca-9,12-dienoate (NHS-LA), was analysed using ^1^H nuclear magnetic resonance (NMR) spectroscopy (Fig. S10A) and mass spectrometry (MS). MS: [M+Na]^+^ 400.5 (calculated), 400.5 (found).

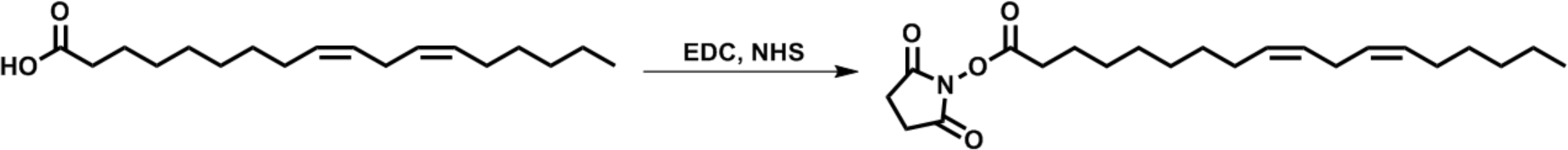

NHS-LA was left O/N at room temperature (RT) to react with L-azidolysine hydrochloride in anhydrous DMF, and produce azide-LA.

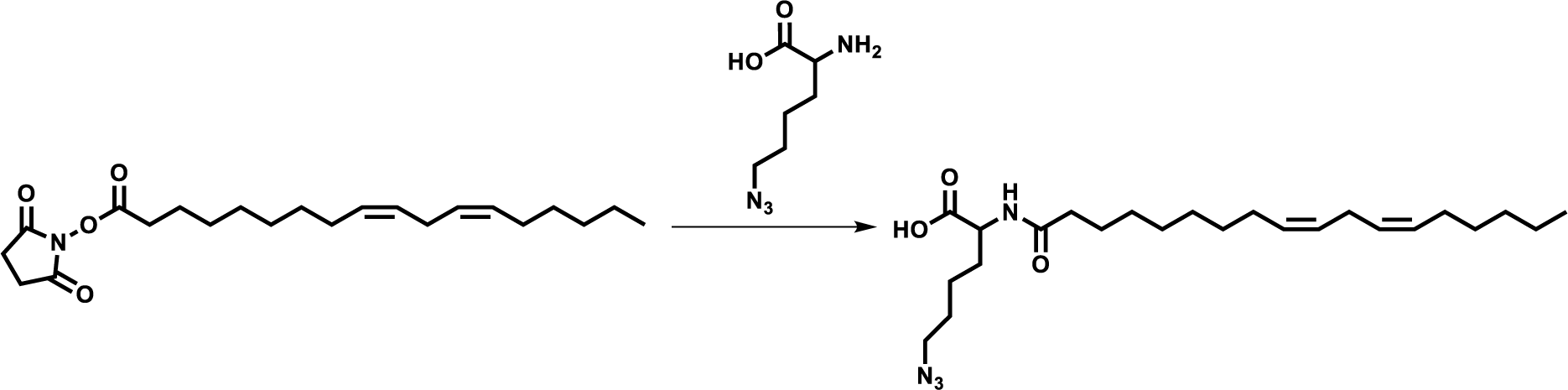

DMF was then removed, the reaction mixture precipitated in water, and dried under vacuum to obtain a clear oil. The composition of the oil was confirmed as being a mixture of azide-LA and unmodified LA (20% and 80%, respectively) based on ^1^H NMR (Fig. S10B) and MS data. MS: [LA-H]^-^ 279.5 (calculated), 279.2 (found), [M-H]^-^ 433.3 (calculated), 433.6 (found).

### Binding assays with azide-LA and click chemistry

*S. aureus* USA300 JE2 WT, Δ*essC*, and Δ*esxC*, grown to OD1, were treated with 10 µM azide-LA for 15 min at 37°C with shaking. The samples were then centrifuged, and the bacterial pellets resuspended in PBS supplemented with 4 µg/mL Click-iT™ Alexa Fluor™ 488 sDIBO alkyne (Life Technologies LTD, UK). After incubation at 25°C for 1h with shaking, bacteria were washed with PBS, and binding to azide-LA was quantified by measuring fluorescence using a FLUOstar OMEGA plate reader (BMG Labtech, UK). The samples imaged with a microscope were additionally stained with 3 µM propidium iodide, following click chemistry. Bacteria stained with Click-iT™ Alexa Fluor™ 488 sDIBO alkyne and 3 µM propidium iodide were immobilized on agarose-covered glass slides, and viewed with a Leica DMi8 widefield microscope (Leica Microsystems LTD, UK). Images were analysed with the ImageJ processing package Fiji (Schindelin *et al*., 2012).

### Live/Dead Staining

Bacteria grown to OD_600_ of 1.0, were treated with 80 μM linoleic acid for 15 min at 37°C with shaking. The samples were then centrifuged, and the bacterial pellets resuspended in PBS and supplemented with a 1:1 ratio of 2X LIVE/DEAD solution (6 μM SYTO-9 stain and 30 μM propidium iodide) from LIVE/DEAD® BacLight kit (Invitrogen). After incubation in the dark for 15 min, bacteria were washed with PBS, spotted on to agarose pads and imaged using a Leica DMi8 widefield microscope (Leica Microsystems, UK). Acquired images were analysed with the ImageJ processing package, Fiji.

### Lipid extraction and analyses

Lipids were extracted from bacterial cultures as described elsewhere (Smith *et al*., 2019). Briefly, bacteria were grown to OD1 in TSB or TSB supplemented with 10 µM LA, centrifuged in a 2 mL glass Chromacol vial (Thermo Scientific), and resuspended in 0.5 mL MS grade methanol (Sigma-Aldrich). MS grade chloroform was then used to extract lipids. The extracted lipids were dried under nitrogen gas with a Techne sample concentrator (Staffordshire, UK), and the lipid pellets resuspended in 1 mL acetonitrile. The samples were then analysed by LC-MS with a Dionex 3400RS HPLC coupled to an amaZon SL quadrupole ion trap mass spectrometer (Bruker Scientific) via an electrospray ionisation interface. Both positive and negative ionisation modes were used for sample analyses. The Bruker Compass software package was utilized for data analyses, using DataAnalysis for peak identification and characterization of lipid class, and QuantAnalysis for quantification of the relative abundance of distinct PG species to total PG species.

### Cell shaving for surface proteome analysis

*S. aureus* USA300 JE2 WT grown to OD1 and Δ*esxC* were washed three times before being treated with Proteomics grade trypsin from porcine pancreas (Sigma-Aldrich, UK) for 15 min as described (Solis *et al*., 2014). The samples were then centrifuged at 1000 × *g* for 15 min, and the bacterial pellets discarded while supernatants were filtered through a 0.2 µM filter. The freshly prepared peptides were frozen (−20°C) until 2 additional, independent biological replicates per strain were prepared.

### Cellular proteomics

*S. aureus* strains were grown O/N at 37°C on tryptic soy agar plates. The next day, single colonies were used to inoculate 10 mL plain TSB or TSB with 10 µM LA. Cultures were grown at 37°C with 180-rpm shaking until an OD600 of 3.2 ± 0.2 was reached. The bacteria were then centrifuged, washed with PBS, and resuspended in lysis buffer (PBS, 250 mM sucrose, 1 mM EDTA, and 50 µg/mL lysostaphin) supplemented with cOmplete^™^, mini, EDTA-free protease inhibitor cocktail (Sigma-Aldrich, UK). After 15 min incubation at 37°C, cells were lysed mechanically with silica spheres (Lysing Matrix B, Fischer Scientific, UK) in a fast-prep shaker as described previously (Mielich-Suss *et al*., 2017). Samples were then centrifuged, and the supernatants transferred to fresh tubes, where proteins were reduced and alkylated for 20 min at 70°C with 10 mM TCEP (tris(2-carboxyethyl)phosphine) and 40 mM CAA (2-chloroacetamide), respectively. Next, the solvent was exchanged first to 8M urea buffer then to 50 mM ammonium bicarbonate (ABC). Proteins were digested O/N at 37°C with mass spectrometry grade lysyl endopeptidase LysC and sequencing grade modified trypsin (Promega LTD, UK).

### Preparation of culture supernatant for proteomics analysis

*S. aureus* strains were grown to an OD600 of 3 in TSB. After centrifugation of cultures, supernatants were sterile-filtered and incubated at 4°C overnight with 10% trichloroacetic acid and 50 µM sodium deoxycholate in presence of one cOmplete™, Mini, EDTA-free Protease Inhibitor Cocktail tablet (Sigma-Aldrich, UK). The precipitated proteins (10 000 × *g* at 4°C for 15 min) were gently washed with acetone and dry at RT for 10 min. After denaturation, proteins were run on a gel until all the proteins had moved from the stacking gel to the resolving gel. The gel was then stained with Instant*Blue*^TM^ (Sigma-Aldrich, UK) for 3 h, after which the protein bands were excised and diced. Proteins were in-gel digested with trypsin as recently described(Goodman *et al*., 2018). Briefly, proteins were reduced and alkylated for 5 min at 70°C with 10 mM TCEP and 40 mM CAA, respectively. Tryptic digestion was carried out at O/N at 37°C in 50 mM ABC.

### Label-free protein quantification

Peptides prepared for proteome analyses were desalted and concentrated with a C18 cartridge in 40 µL MS buffer (2% acetonitrile plus 0.1% trifluoroacetic acid). For each sample, 20 µL were analysed by nanoLC-ESI-MS/MS using the Ultimate 3000/Orbitrap Fusion instrumentation (Thermo Scientific), and a 90-minute LC separation on a 50 cm column. The data were used to interrogate the Uniprot *Staphylococcus aureus* USA300 database UP000001939, and the common contaminant database from MaxQuant (Cox *et al*., 2014). MaxQuant software was used for protein identification and quantification using default settings. Intensities were log_2_-tansformed with the Perseus software, and proteins with one or no valid value for every sample in triplicate were filtered. For surfome data, the removeBatchEffect function of the limma R package (Ritchie *et al*., 2015) was used to remove differences accounting for variation in shaving efficiency done on three different days for all the biological replicates. Missing values in cellular proteomics data were imputed on R. Specifically, for each sample, the imputed value was either the lowest intensity across all samples if at least two biological replicates had missing values or the average of two valid values if only one was missing.

### ROS measurement

Chemical hydrolysis of 2,7-dichlorofluorescein diacetate (Sigma-Aldrich) was performed to acquire a final dichlorofluorescein (DCF) reagent yield of 50 μM. Briefly, 0.5 ml of 5 mM DCF-DA (dissolved in 100% ethanol), was reacted with 2 ml of 0.1 N NaOH at RT for 30 min. The reaction was stopped using 7.5 mL 10X PBS pH 7.4 (without calcium and magnesium; Gibco). Bacteria grown to OD_600_ of 1.0 were treated with 10µM linoleic acid or left untreated for 15 min at 37°C shaking and 1 mL of this culture was centrifuged. Cells were resuspended in 100 µL of DCF reagent and incubated for 40 min in the dark at RT. Fluorescence was measured using a FLUOstar OMEGA plate reader (BMG Labtech, UK) at an excitation wavelength of 482 nm and emission wavelength of 520 nm.

### Data availability statement

The mass spectrometry proteomics data have been deposited to the ProteomeXchange Consortium via the PRIDE (Perez-Riverol *et al*., 2019) partner repository with the dataset identifier PXD013081 and 10.6019/PXD013081. Cellular proteomic and surface proteome samples are labelled MS18-193 and MS17-185, respectively.

### Statistical analyses

Except for the proteomics results, the statistical tests were performed with GraphPad Prism 8.0 as indicated in the Figure legends, with *P* values < 0.05 considered significant. A paired two-tailed Student’s t-test or a paired Mann-Whitney U test was used for pairwise comparisons. An ordinary one-way analysis of variance (ANOVA) with Dunnett’s multiple comparisons test or a Kruskal-Wallis test with Dunn’s multiple comparisons test was applied to data form three or more groups. The fold changes and *P* values of the proteomics data were calculated with the R package limma (Ritchie *et al*., 2015), with USA300 JE2 WT or bacteria grown without LA as references. These fold changes and *P* values were used by the R package piano (Varemo *et al*., 2013) to compute the enrichment of gene ontology (GO) terms.

## Supporting information

Suppmlmentary tables

supplementary proteomics data

## Acknowledgments

We thank Professor Tracy Palmer (Newcastle University) and Professor Olaf Scheewind (University of Chicago) for providing us *S. aureus* strains and reagents. We thank GSK, Siena, Italy for providing the *esxA*, *esxB* mutant strains used in this study. We acknowledge the contribution of the Proteomics Research Technology Platform, University of Warwick. The authors have no conflict of interest to declare.

---

**Fig. S1. USA300 JE2 WT and Δ*esxC* strains display similar growth rates.** WT and Δ*esxC* were grown in TSB, and OD_600_ monitored with a Novaspec^®^ Pro spectrophotometer. Data shown are means of three independent experiments, and the error bars indicate the standard errors of the mean.

**Fig. S2. EsxC associates with *S. aureus* membrane**. Immunoblot analysis of cell membrane (CM) or cell wall (CW) fractions of WT (USA300), Δ*essC*, and Δ*esxC* with anti-EsxC sera or anti-PBP2a antibodies (loading control).

**Fig. S3. Volcano plot of the quantitative proteomic analysis of surface proteins in Δ*esxC* compared to WT.** The relative abundance of each protein (log_2_ fold change, X-axis) and its statistical significance (*P* value, Y-axis) are shown in the graph. Proteins decreased by more than half in Δ*esxC* (log_2_ fold change < −1 and *P* value < 0.05) are shown in green.

**Fig. S4. *S. aureus* growth inhibition by arachidonic acid is increased in T7SS mutants.** *S. aureus* WT USA300, Δ*essC*, and Δ*esxC* were grown in TSB or TSB supplemented with 80 µM arachidonic acid (AA). Means ± standard error of the mean (SEM) are shown, *n* = 3.

**Fig. S5. T7SS substrates contribute to resistance to linoleic acid toxicity.**

A. *S. aureus* USA300 wild-type (WT) and USA300 *esxA* (Δ*esxA*) or *esxB* (Δ*esxB*) deletion mutants were grown in TSB or TSB supplemented with 80 µM linoleic (LA) or stearic acid (SA).

B. *S. aureus* Newman WT and Newman *esxA* (Δ*esxA*) or *esxB* (Δ*esxB*) deletion mutants were grown similarly in TSB or TSB + 40 µM LA or SA.

C. Growth curves as described in (A) were done with RN6390 wild-type (WT) and RN6390 *essC* (Δ*essC*) or *esxC* (Δ*esxC*) deletion mutants. Data shown in (A), (B), and (C) are representative of at least three independent experiments.

**Fig. S6. USA300 JE2 WT and T7SS mutants display similar lipids.**

Representative HPLC chromatograms of the indicated bacteria grown in TSB (A) or in TSB supplemented with LA (B), in negative ionisation mode. Phosphatidylglycerol (PG) is highlighted.

C. Representative HPLC chromatograms of native PG species of Δ*essC* grown in TSB (top panel) or in TSB supplemented with LA (bottom panel), in negative ionisation mode.

**Fig. S7. LA (C18:2) is elongated and incorporated into *S. aureus* phosphatidylglycerol (PG) species.** A-C. Representative mass spectrometry fragmentation spectra for PG species containing unsaturated fatty acids, in negative ionisation mode.

A. PG species with mass 731 m/z, containing C18:2 fatty acid (279 m/z).

B. PG species 759 m/z, containing C20:2 fatty acid (307 m/z)

C. PG species 787 m/z, containing C22:2 fatty acid (335 m/z).

D. and E. Relative quantification of the indicated PG species containing an unsaturated fatty acid in WT, Δ*essC* and Δ*esxC*. C20:2-(D) and C22:2-containing PG species (E) are presented as ratios of total PG species. Data shown are the means and error bars represent SD of three independent experiments.

**Fig. S8. USA300 JE2 WT and Δ*esxC* growth similarly in presence of sub-inhibitory amounts of linoleic acid.** *S. aureus* WT USA300 and Δ*esxC* were grown in TSB or TSB supplemented with 40 µM linoleic acid (LA). Means ± SEM are shown, *n* = 3.

**Fig. S9. Principal component analysis (PCA) of the *S. aureus* cellular proteomic profiles.** PCA was performed on all the identified proteins of USA300 JE2 WT and T7SS mutants grown in TSB (untreated) or TSB + LA (LA-treated). Each dot represents a biological replicate.

**Fig. S10. ^1^H NMR spectra of NHS-LA (A) and azide-LA (B) in CDCl_3_.** Both spectra were recorded on a Bruker Advance 300 spectrometer (300 MHz) at 27 °C. The letters indicate the chemical shift δ (in parts per million, ppm) of the protons in each molecule.

**Dataset S1. Differentially abundant proteins in WT USA300 JE2 and T7SS mutants in response to linoleic acid.**

