## Supplementary figures and images for "The type VII secretion system protects *Staphylococcus aureus* against antimicrobial host fatty acids"

### Supplementary figures

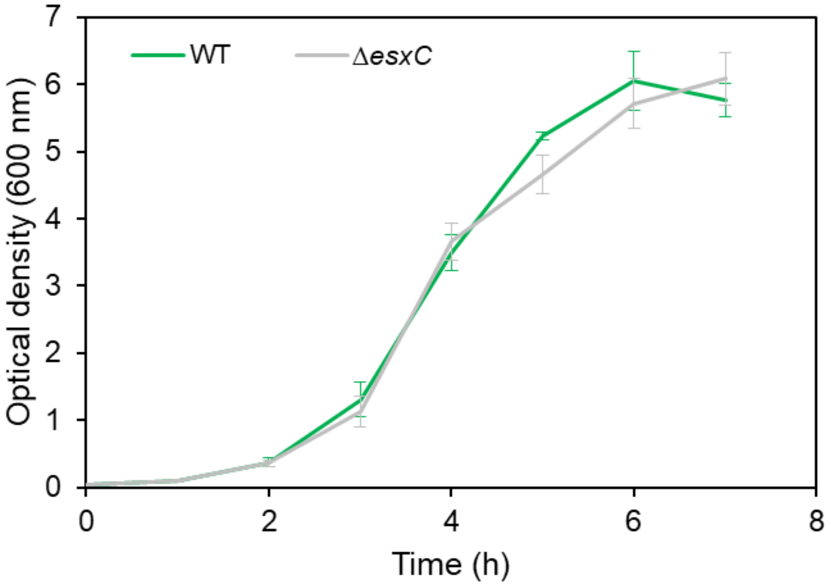


**Fig S1**

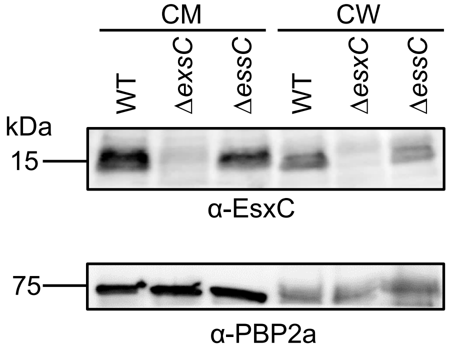


**Fig S2**

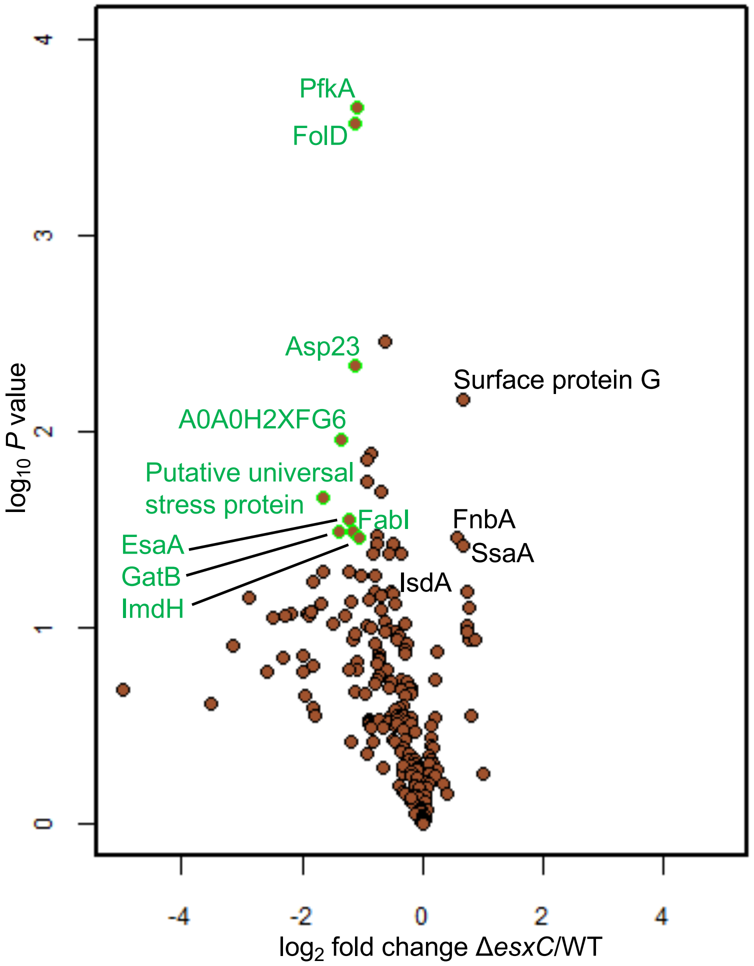

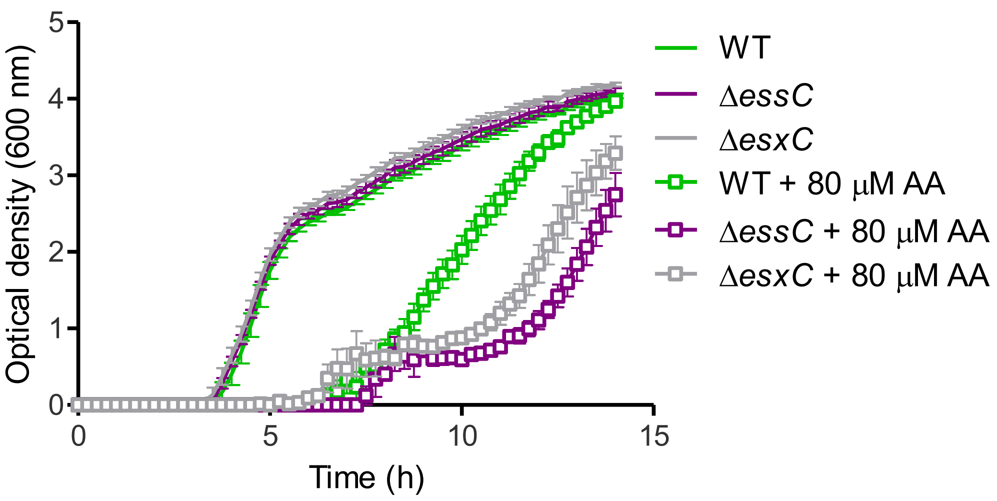

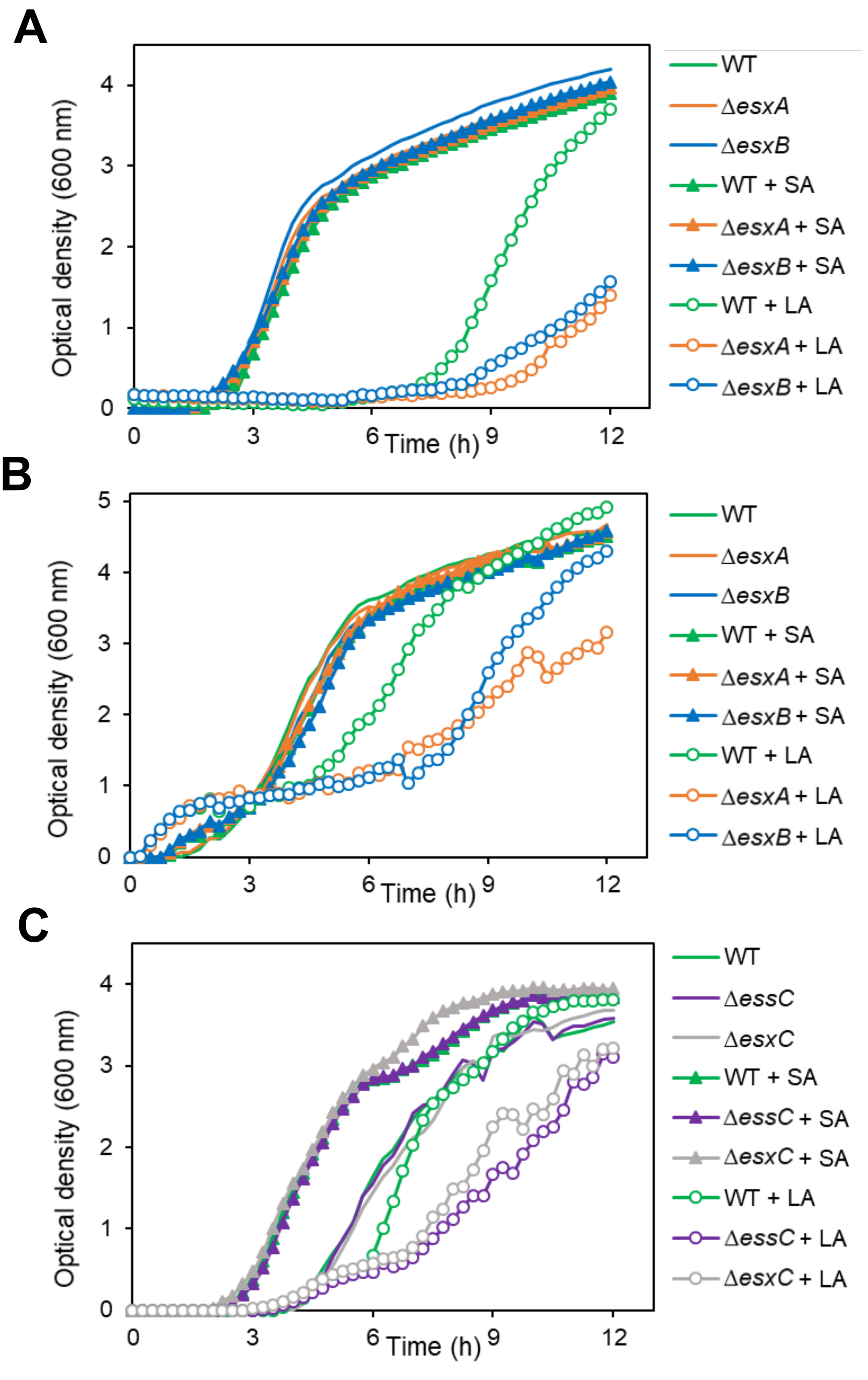

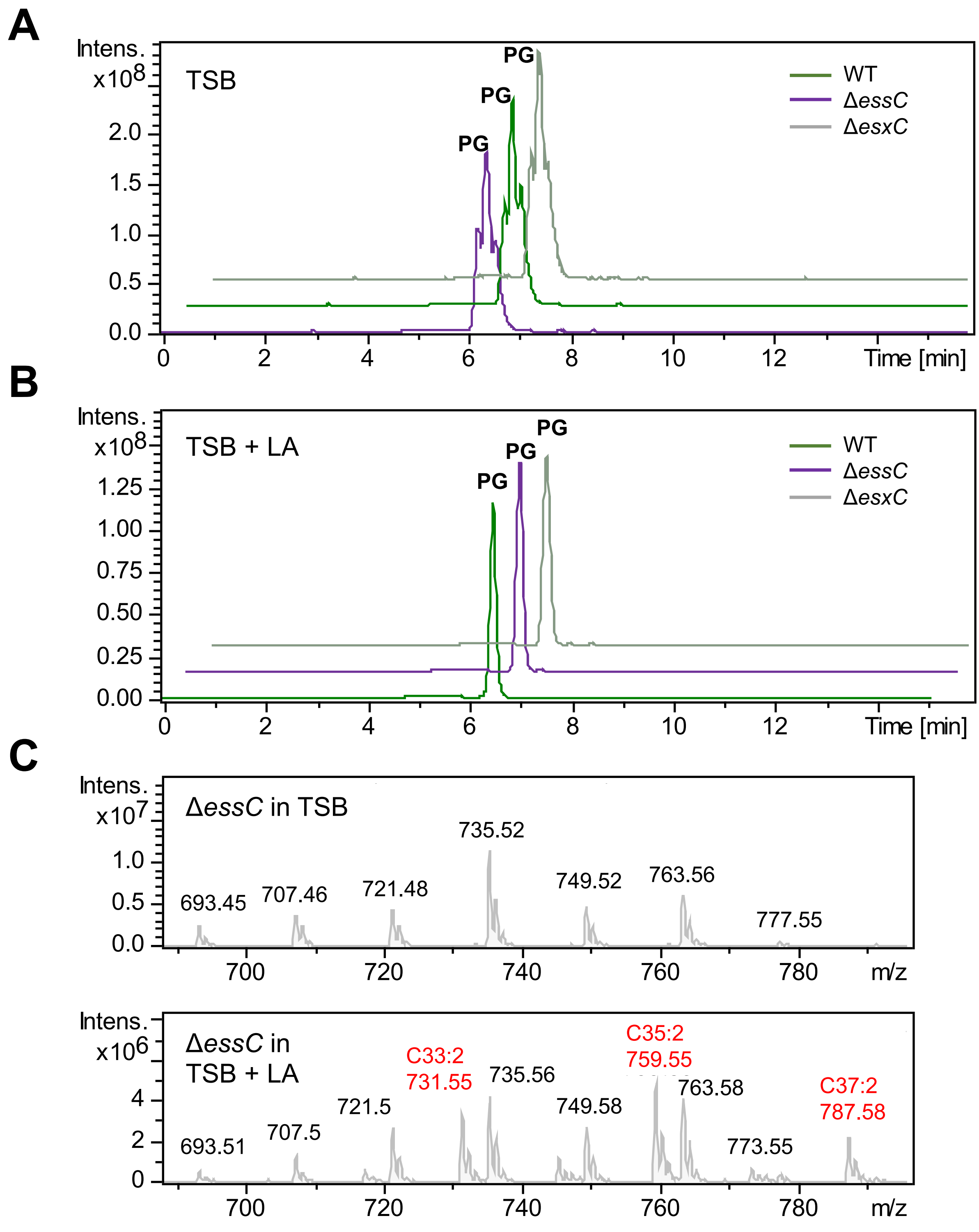

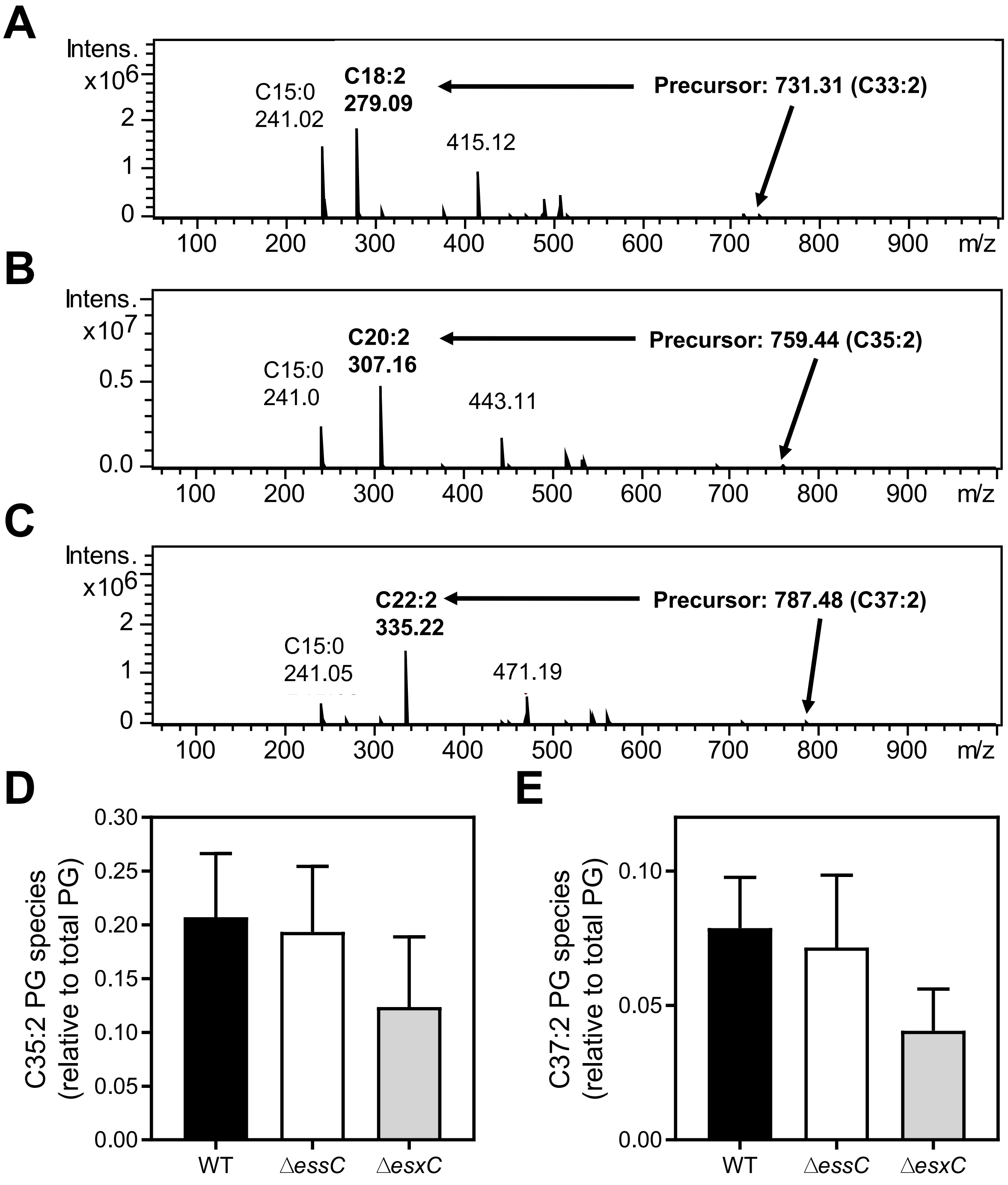

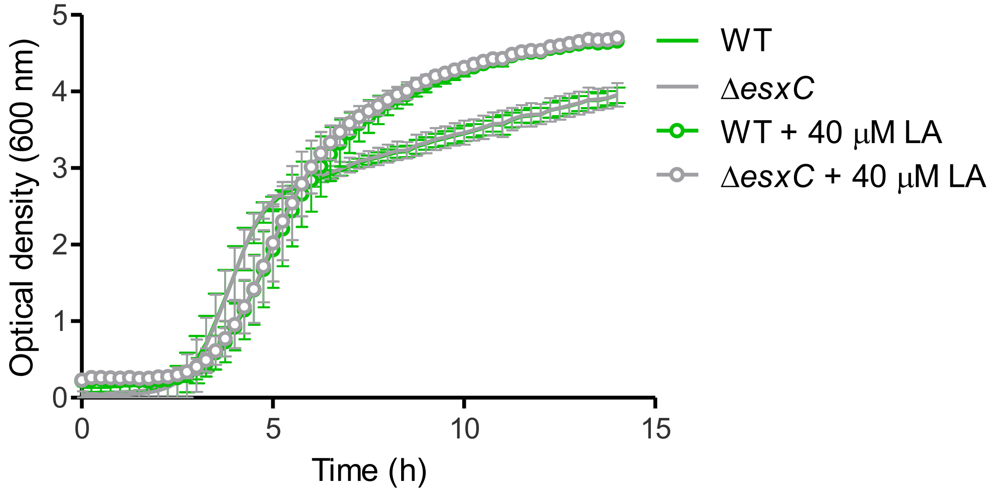

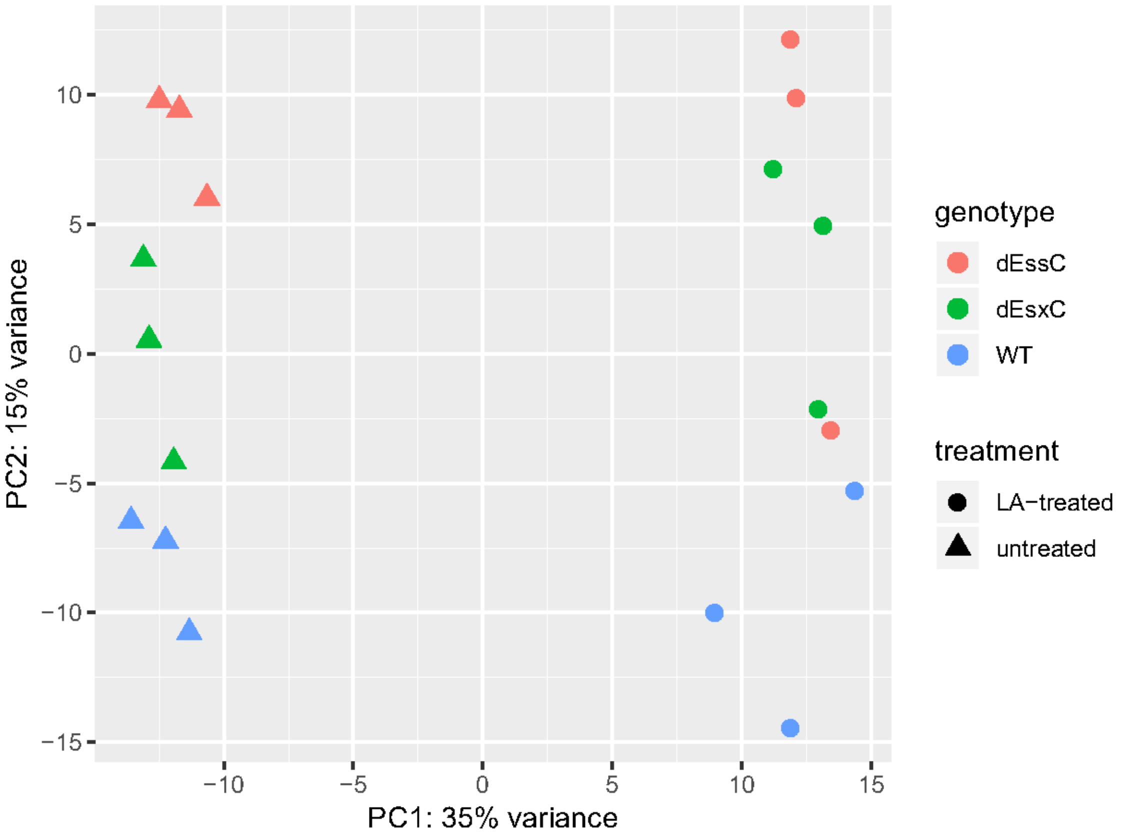

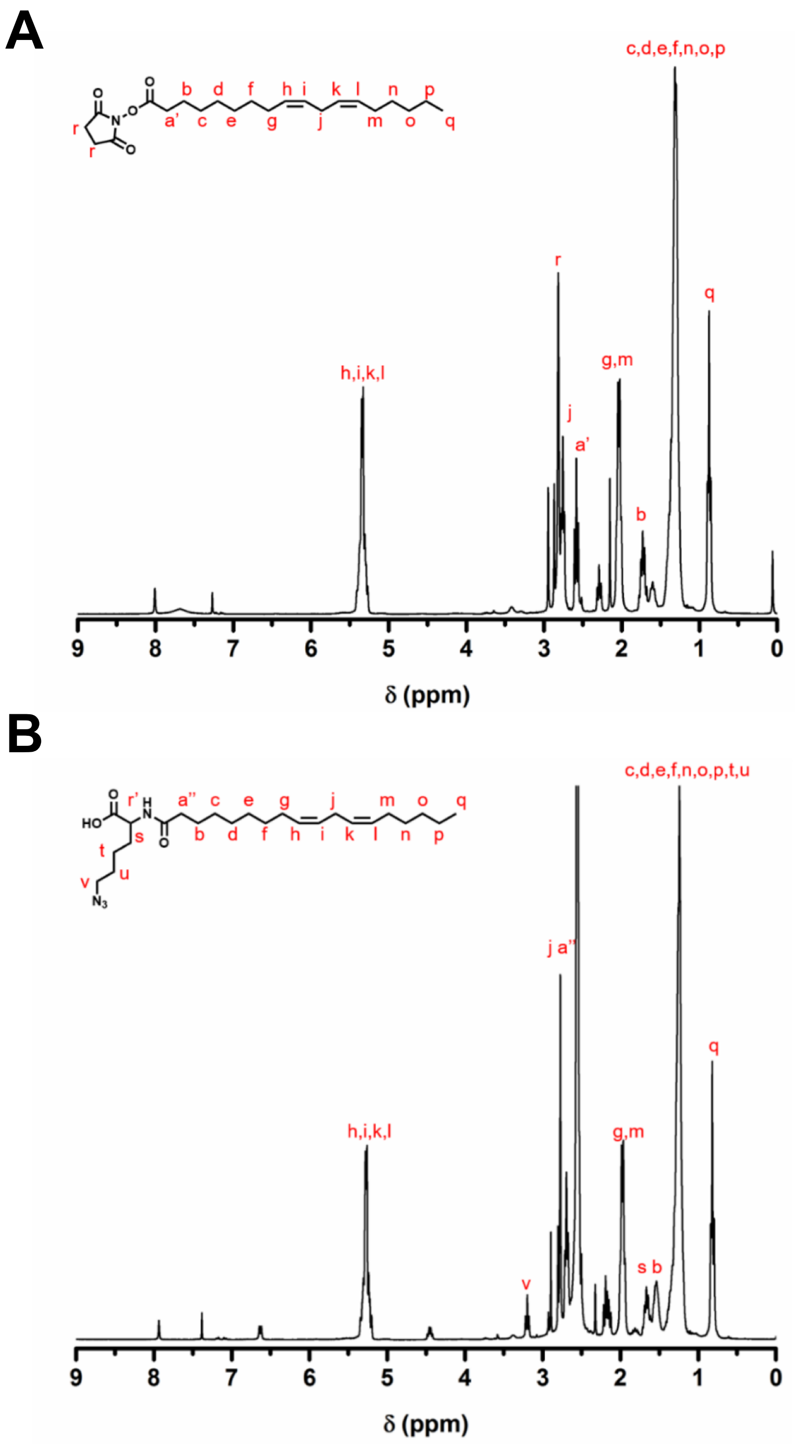


Fig S10
