## Supplementary material for "The type VII secretion system protects *Staphylococcus aureus* against antimicrobial host fatty acids": Suppmlmentary tables

**Table S1. Secreted proteins significantly changed in Δ*esxC* and Δ*essC* mutants relative to WT USA300 JE2**

| | Uniprot ID | $log_2$ fold change | Adjusted *P* value | Description |
|---|---|---|---|---|
| USA300 JE2 Δ*esxC* vs USA300 JE2 WT | A0A0H2XIV9 | -4.27 | 7.11E-10 | T7SS protein EsxD |
| | A0A0H2XHT7 | -2.78 | 7.96E-09 | Phosphocarrier protein HPr |
| | A0A0H2XH98 | -1.75 | 1.34E-07 | Hydrolase, MutT/nudix family |
| | A0A0H2XIK2 | -1.71 | 1.34E-07 | T7SS protein EsxC |
| | A0A0H2XI99 | -1.45 | 0.003045 | T7SS protein EsxA |
| | A0A0H2XFA5 | -1.28 | 4.03E-05 | Putative GTP-binding protein |
| | A0A0H2XH08 | -0.76 | 2.77E-05 | Accessory gene regulator protein A AgrA |
| | Q2FDS6 | -0.75 | 2.77E-05 | Uncharacterized hydrolase |
| | A0A0H2XH39 | 0.71 | 3.61E-05 | Cell division protein FtsL |
| | A0A0H2XEF1 | 1.55 | 3.40E-06 | 2-oxoisovalerate dehydrogenase |
| | Q2FEF1 | 1.63 | 0.000836 | Isopentenyl-diphosphate delta-isomerase |
| | A0A0H2XKB6 | 1.90 | 0.00012 | Uncharacterized protein |
| | A0A0H2XH53 | 2.58 | 0.042723 | Uncharacterized protein |
| USA300 JE2 Δ*essC* vs USA300 JE2 WT | A0A0H2XI99 | -6.59779 | 1.63E-07 | T7SS protein EsxA |
| | A0A0H2XIV9 | -4.27062 | 7.11E-10 | T7SS protein EsxD |
| | A0A0H2XHT7 | -2.77719 | 7.96E-09 | Phosphocarrier protein HPr |
| | A0A0H2XKB6 | -2.39392 | 2.34E-05 | Uncharacterized protein |
| | A0A0H2XH98 | -1.74886 | 1.34E-07 | Hydrolase, MutT/nudix family |
| | A0A0H2XIK2 | -1.70544 | 1.34E-07 | T7SS protein EsxC |
| | A0A0H2XJ55 | -1.24877 | 0.000168 | Uncharacterized protein |
| | Q2FHI5 | -1.04906 | 0.015225 | ATP-dependent protease subunit HslV |
| | A0A0H2XII7 | -0.87542 | 0.000163 | Exotoxin |
| | Q2FHT6 | -0.82647 | 0.035546 | Thioredoxin |
| | Q2FH36 | -0.82371 | 0.016379 | Cold shock protein CspA |
| | Q2FDK5 | -0.78489 | 0.015225 | Serine-rich adhesin for platelets |
| | A0A0H2XH08 | -0.76352 | 2.12E-05 | Accessory gene regulator protein A AgrA |
| | Q2FDS6 | -0.74909 | 2.15E-05 | Uncharacterized hydrolase |
| | Q2FJ70 | -0.74787 | 0.015871 | 3-hexulose-6-phosphate synthase |
| | Q2FGE2 | -0.61344 | 0.010461 | Protein GrpE |
| | Q2FEZ4 | -0.51905 | 0.039832 | S-ribosylhomocysteine lyase |
| | A0A0H2XG16 | -0.5063 | 0.021123 | Clumping factor A |
| | A0A0H2XGZ2 | -0.5033 | 0.010741 | Uncharacterized protein |
| | Q2FF55 | 0.628649 | 0.021219 | Alanine racemase |
| | Q2FFT2 | 0.777989 | 0.026729 | Serine protease |
| | A0A0H2XEF1 | 1.394088 | 5.08E-06 | 2-oxoisovalerate dehydrogenase |
| | A0A0H2XHA0 | 1.470262 | 0.041559 | Uncharacterized protein |
| | A0A0H2XG38 | 1.580427 | 1.63E-07 | Uncharacterized protein |
| | Q2FEF1 | 1.633554 | 0.000707 | Isopentenyl-diphosphate delta-isomerase |

**Table S2. Surface proteins significantly changed in Δ*esxC* mutant relative to WT USA300 JE2.**

| Uniprot ID | $log_2$ FC | *P* Value | Description | Localization |
|---|---|---|---|---|
| A0A0H2XJV9 | 0.7 | 0.0069 | Surface protein G | Cell wall |
| A0A0H2XHF6 | 0.7 | 0.0383 | Secretory antigen SsaA | Extracellular |
| Q2FE03 | 0.6 | 0.0350 | Fibronectin-binding protein A FnbA | Cell wall |
| Q2FG28 | -1.7 | 0.0217 | Putative universal stress protein | Cytoplasm |
| A0A0H2XFG6 | -1.4 | 0.0110 | Uncharacterized secreted protein | Extracellular |
| Q2FFJ6 | -1.4 | 0.0326 | Amidotransferase subunit B GatB | Cytoplasm |
| A0A0H2XFP1 | -1.2 | 0.0281 | ESAT-6 secretion accessory factor EsaA | Membrane |
| A0A0H2XGD7 | -1.1 | 0.0002 | ATP-dependent 6-phosphofructokinase PfkA | Cytoplasm |
| Q2FI15 | -1.1 | 0.0003 | Bifunctional protein FolD | Cytoplasm |
| Q2FEV0 | -1.1 | 0.0046 | Alkaline shock protein 23 | Cytoplasm |
| A0A0H2XIJ1 | -1.1 | 0.0322 | Enoyl-[acyl-carrier-protein] reductase [NADPH] FabI | Cytoplasm |
| Q2FJM6 | -1.0 | 0.0345 | Inosine-5-monophosphate dehydrogenase | Cytoplasm |
| A0A0H2XDE4 | -0.9 | 0.0130 | Transcriptional regulator MgrA | Cytoplasm |
| Q2FJM5 | -0.9 | 0.0140 | GMP synthase [glutamine-hydrolyzing] | Cytoplasm |
| Q2FFV6 | -0.9 | 0.0182 | S-adenosylmethionine synthase MetK | Cytoplasm |
| A0A0H2XGP3 | -0.8 | 0.0339 | UDP-N-acetylglucosamine 1-carboxyvinyltransferase MurA | Cytoplasm |
| Q2FJA0 | -0.8 | 0.0421 | 50S ribosomal protein L7/L12 | Cytoplasm |
| A0A0H2XH82 | -0.7 | 0.0202 | Putative lipoprotein | Membrane |
| Q2FJE0 | -0.7 | 0.0378 | 50S ribosomal protein L25 | Cytoplasm |
| A0A0H2XFW2 | -0.6 | 0.0035 | Universal stress family protein | Cytoplasm |
| Q2FG40 | -0.5 | 0.0373 | Pyruvate kinase | Cytoplasm |
| Q2FHV1 | -0.5 | 0.0423 | Iron-regulated surface determinant protein A IsdA | Cell wall |
| Q2FID4 | -0.4 | 0.0421 | NADH dehydrogenase-like protein | Cytoplasm |

**Table S3. Strains and plasmids used in this study.**

| Strain or plasmid | Description | Source or reference |
|---|---|---|
| ***Staphylococcus aureus*** | | |
| USA300 LAC | Community-acquired MRSA (CA-MRSA) | Olaf Schneewind |
| LAC Δ*esxA* | *S. aureus* USA300 LAC defective for EsxA | (Korea *et al.*, 2014) |
| LAC Δ*esxB* | *S. aureus* USA300 LAC defective for EsxB | (Korea *et al.*, 2014) |
| USA300 LAC JE2 | Plasmid-cured USA300 LAC | BEI Resources (Fey *et al.*, 2013) |
| JE2 Δ*esxC* | *S. aureus* USA300 LAC JE2 defective for EsxC | This study |
| JE2 Δ*essC* | *S. aureus* USA300 LAC JE2 defective for EssC | This study |
| Newman | Methicillin-sensitive *Staphylococcus aureus* | Olaf Schneewind |
| Newman Δ*esxA* | *S. aureus* Newman defective for EsxA | (Korea *et al.*, 2014) |
| Newman Δ*esxB* | *S. aureus* Newman defective for EsxB | (Korea *et al.*, 2014) |
| RN6390 | NCTC8325 derivative, Δ*rbsU*, Δ*tcaR*, cured of φ11, φ12, and φ13 | Tracy Palmer (Kneuper *et al.*, 2014) |
| RN6390 Δ*esxC* | *S. aureus* RN6390 defective for EsxC | Tracy Palmer (Kneuper *et al.*, 2014) |
| RN6390 Δ*essC* | *S. aureus* RN6390 defective for EssC | Tracy Palmer (Kneuper *et al.*, 2014) |
| RN4220 | *S. aureus* restriction negative, cloning tool | BEI Resources (NARSA) |
| **Plasmids** | | |
| pKORI | Temperature-sensitive allelic exchange vector | Olaf Schneewind (Bae & Schneewind, 2006) |
| pKORIΔ*essC* | pKORI used to generate *essC* mutant | This study |
| pKORIΔ*esxC* | pKORI used to generate *esxC* mutant | This study |
| pOS1 | Insertless vector for genetic complementation | Olaf Schneewind |
| pOS1CK | pOS1 with P1 constitutive promoter of *sarA* | (Korea *et al.*, 2014) |
| pOS1-*esxA* | *esxA* complementation vector | (Korea *et al.*, 2014) |
| pOS1-*esxC* | *esxC* complementation vector | This study |

**Table S4. Primers used in this study.**

| Primer | Sequence (5' – 3') | Product |
|---|---|---|
| AttB1-*esxC* Up-Fwrd | GGGGACAAGTTTGTACAAAAAAGCAGGCTGAGCTAACGCTATGAAAACACC | AttB1-Δ*esxC*-AttB2 |
| *esxC*-up-Rev-soeing | ACCCATATCTTCACCTCAATAAACATACCTCCCTCCTATTT | |
| *esxC*-down-Fwd | TATTGAGGTGAAGATATGGGTGG | |
| *esxC*-down-Rev-attB2 | GGGGACCACTTTGTACAAGAAAGCTGGGTCGTCATTACTCCTCTGCTTTA | |
| AttB1-*essC*-up-fwd | GGGGACAAGTTTGTACAAAAAAGCAGGCTGCTACACATTTGTGTTGGCACC | AttB1-Δ*essC*-AttB2 |
| *ess*C-upstream-rev | TGTCTTTGCCTCAGTCCTATAC | |
| *essC*-dpwn-soeing-fwd | GTATAGGACTGAGGCAAAGACACAATGAATTAAATAGGAGGGAGG | |
| *esxC*-down-Rev-attB2 | GGGGACCACTTTGTACAAGAAAGCTGGGTCGTCATTACTCCTCTGCTTTA | |
| *esxC*-RBS-PstI-fwd | GCGCTGCAGTTGAGAGGAGAGAAAATGAATTTTAATGATATTGAAAC | RBS-*esxC* |
| *esxC*-SmaI-rev | GCGGCGCCCGGGTTAATTCATTGCTTTATTAAA | |
